# Luminal progenitors undergo partial epithelial-to-mesenchymal transition at the onset of basal-like breast tumorigenesis

**DOI:** 10.1101/2022.06.08.494710

**Authors:** Camille Landragin, Melissa Saichi, Pacôme Prompsy, Adeline Durand, Jérémy Mesple, Amandine Trouchet, Marisa Faraldo, Hélène Salmon, Céline Vallot

**Affiliations:** CNRS UMR3244, Institut Curie, PSL University, Paris, France; Translational Research Department, Institut Curie, PSL University, Paris, France; INSERM U932, Institut Curie, PSL University, Paris, France; Single Cell Initiative, Institut Curie, PSL University, Paris, France; CNRS UMR3215, Institut Curie, PSL University, Paris, France; INSERM U934, Institut Curie, PSL University, Paris, France

## Abstract

Defects in double-strand repair mechanisms - both through germline or somatic inactivation of repair genes - is a hallmark of basal-like breast cancers. In this genetically-unstable context, a recurrent shift in cell identity occurs within the mammary epithelium. Basal-like tumors have indeed been proposed to originate from luminal progenitor (LP) cells yet tumor-initiating events remain poorly understood. Here, we map state transitions at the onset of basal-like tumorigenesis, using a Brca-1 deficient mouse model launching tumorigenesis in multiple LP cells. Combining single-cell transcriptomics to spatial multiplex imaging, we identify a population of cycling p16-expressing cells, emerging from the luminal progenitor compartment, undergoing partial epithelial-to-mesenchymal transition and losing luminal identity. Pseudo-temporal analyses position these cells as a transitory pre-tumoral state between aberrant Brca1-deficient luminal progenitors and growing tumor cells. In patients, the p16 pre-tumoral signature is found in early stage basal-like tumors, that rarely recur. Concomitant to p16 activation, we show that LP cells undergo an epigenomic crisis attested by the general re-organization of their heterochromatin. They accumulate multiple H3K27me3 micro-foci - reminiscent of the formation of senescence-associated heterochromatin foci (SAHFs) - and lose their inactive X (Xi). Both p16 activation and heterochromatin reorganization are hallmarks of human basal-like breast tumors; we propose that these events occur during initial LP transformation and are scars of an initial transitory senescent-like state.

## INTRODUCTION

Triple-negative breast cancer (TNBC) refers to a subgroup of aggressive breast cancers defined by the lack of estrogen receptor (ER), progesterone receptor (PR), and human epidermal growth factor receptor 2 (HER2) accounting for 15–20% of all breast tumors (Onitilo et al. 2009). Along with transcriptional heterogeneity, TNBC is characterized by complex genomes, dictated by high genetic instability and complex patterns of copy number alterations and chromosomal rearrangements (Gao et al. 2016; Engebraaten, Vollan, and Børresen-Dale 2013). Defects in double-stranded DNA repair mechanisms are indeed characteristic of TNBC, as a result of either germline or somatic mutations in BRCA1/2 and other genes involved in DNA repair (Timms et al. 2014; Stefansson et al. 2011). In this genetically unstable context, there is a chaotic de-structuration of the mammary gland, with recurrent loss of proper cell identity. Part of these cancers harbor basal-like phenotypes, expressing an incomplete set of basal markers but with high intra-tumor heterogeneity (Marra et al. 2020; Bianchini et al. 2016). Interestingly, BRCA1-deficient tumors are suspected to originate from HDR-deficient luminal progenitor cells of the gland, implicating a recurrent switch or loss in cell identity during tumorigenesis (Molyneux et al. 2010; E. Lim et al. 2009). Recent data indicate that Brca1-deficiency in the mammary gland induces aberrant alveolar differentiation of luminal progenitors, suggesting early phenotypic defects in the mammary gland of a Brca1-deficient individual (Bach et al. 2021). However, the tumor-initiating events leading to the emergence of tumor cells per se remain unknown.

Studying early steps of tumorigenesis is not feasible solely based on human tumor samples, which are complex stacks of molecular alterations acquired over time. Animal models enable the isolation of a continuum of states from normal to pathologic gland to precisely map the evolution of the physiological mammary gland towards tumorigenesis. In the case of basal-like breast cancers, models with Brca1/Trp53 deficiency in luminal progenitors have been shown to mimic formation of human basal-like breast cancers (Selbert et al. 1998; Molyneux et al. 2010). *TP53* mutations remain the most common genetic alteration in basal-like cancers (85%, (Cancer Genome Atlas Network 2012)). In *BRCA1*-germline carriers, *TP53* mutation was actually shown to be among the earliest events in tumor formation (Martins et al. 2012). In this context, a mouse model with conditional deletion of *Trp53* and *Brca1* in the luminal compartment of the mammary gland appears as an apropos model to catch the rare transforming events leading an HDR-deficient luminal progenitor to tumorigenesis. In contrast to humans, where these events are extremely rare, the deletion in the mouse of these genes in multiple cells of the mammary glands greatly enhances our ability to detect the transitioning states from aberrant luminal progenitor to basal-like breast cancer phenotype.

Here, using single-cell transcriptomics and multiplex imaging in a Blg-Cre Trp53^Fl/Fl^ and Brca1^Fl/Fl^ mouse model, we mapped steps of Brca1-tumorigenesis *in vivo*, with a focus on epithelial cells to catch rare pre-tumoral epithelial states. We identified an intermediate population of cells, expressing p16, transitioning from luminal progenitor to tumor phenotype, with highly remodeled genomes. These cells are partially switching to a mesenchymal phenotype while retaining their epithelial characteristics and activating angiogenesis. We furthermore discovered that LPs concomitantly undergo a major epigenomic crisis with a disruption of their heterochromatin through the accumulation of multiple heterochromatin foci and loss of their Xi, a hallmark of basal-like breast tumors. Using TCGA data, we further show that the p16-associated pre-tumoral signature is found in basal-like cancers with rare recurrence. We propose that p16 activation and heterochromatin disruption could be scars of an early senescence-like transitory state in the basal-like tumorigenesis process.

## RESULTS

### Monitoring mammary tumorigenesis with scRNA-seq *in vivo*

Virgin Blg-Cre Trp53^fl/fl^ Brca1^fl/fl^ females develop mammary tumors at a median age of 5.2 months (Fig. S1A). These tumors display a complete de-structuration of the mammary gland, with high intra-tumoral heterogeneity: 72% of cells express neither the canonical basal marker Krt5 nor the luminal marker Krt8, 17% are positive for Krt8 only, 0,2% for Krt5 only whereas 11% of cells expressed both markers, revealed by immunofluorescent staining (Fig. S1B). In order to delineate the steps leading to such destructuration, we profiled mammary epithelium from animals at various time points (2.7, 3.2 and 5.2 months, n=12), as well as from three tumors (Fig. 1A, n=15 mice in total). To maximize our chances of identifying tumor-initiating cells, among the n=15 mice, we profiled the mammary epithelium of 2 mice at 5.2 months of age, with no apparent tumor, but belonging to a litter of an animal with a tumor. Among these individuals at 5.2 months of age, we observed multiple lesions - less than 0,5 mm - within the mammary epithelium (Fig. 1A). Part of the collected samples were enriched for epithelial fraction to further increase our chances of identifying rare phenotypic states within the Brca1/Trp53 deficient mammary epithelium (see Methods), on which scRNA-seq was performed using 10X technology.

**Figure 1:**
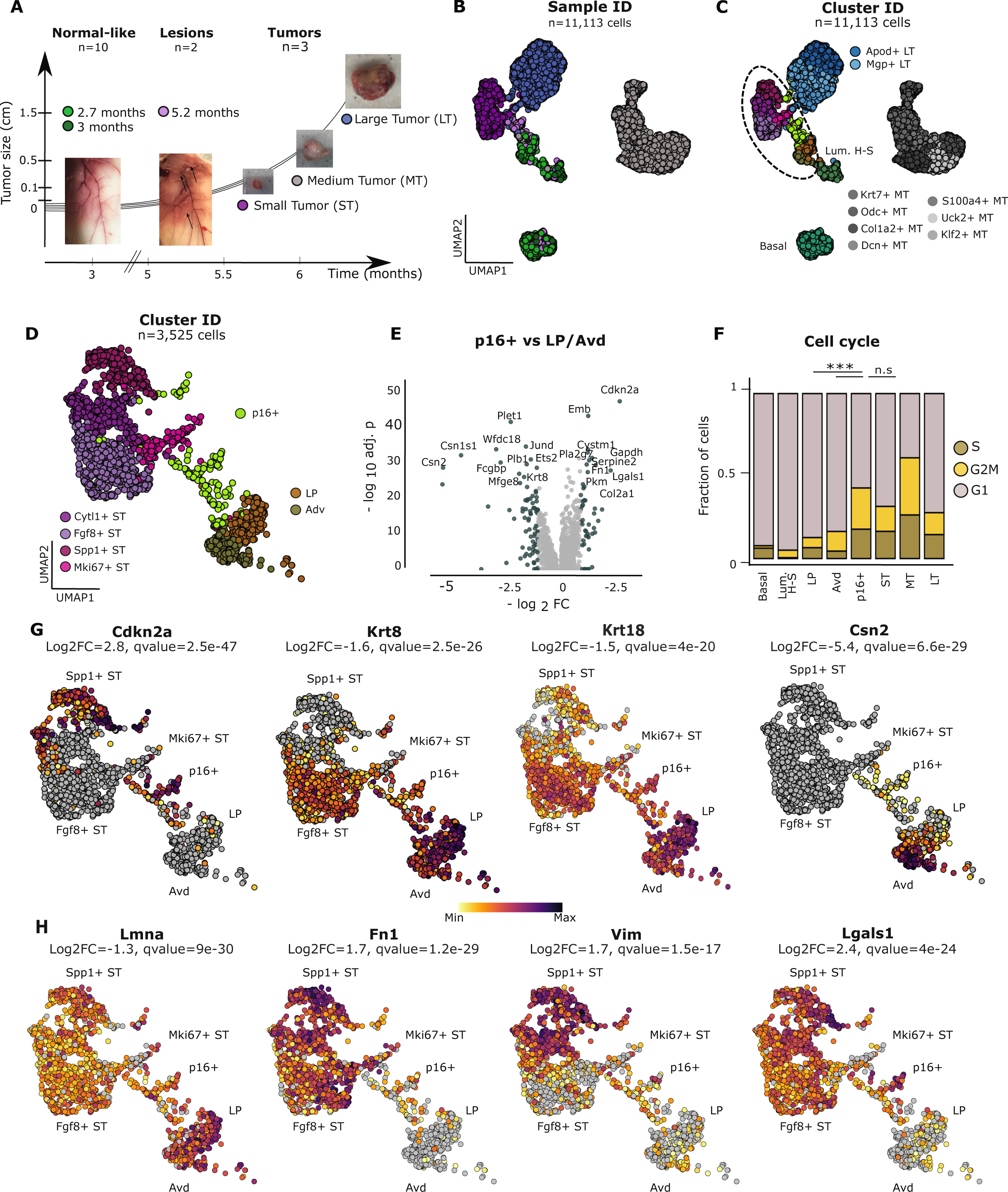
Identification of p16-high cycling cells within lesions of the mammary gland. (A) Schematic representation of the timeframe and histological classification of the in vivo processed samples using scRNAseq 10X technology. (B) UMAP representation of the epithelial cells; each dot represents a cell, and is colored according to the sample of origin. (C) UMAP, colored according to cluster-based annotation. (D) Zoom in on the corresponding UMAP embeddings to the transitional cell clusters (see material and methods). Cells were colored according to cluster-based annotation. (E) Volcano plot representation of the DEGs in p16+ cycling cells compared to LP and Avd, highlighted are top DEG, with an absolute log2_FC > 1.2 and a significant adjusted p-value (<0.05). Xaxis represents the log2 FC and the y-axis represents the -log10(adjusted p-value). (F) Barplot representation of the fraction of cell cycle phases (G1,S or G2/M) inferred for each Major-type annotation; including: Basal, Luminal Hormone-Sensing (H-S), Luminal Progenitors (LP), Alveolar-differentiated (Avd), P16+lesional cells, and grouped cells per tumor size: Small Tumor (ST), Medium tumor (MT) and late tumor (LT). asterisks above LP indicate significance of p16+ versus LP; asterisks above ST indicate significance of p16+ versus ST. *P < 0.05, **P <0.01, ***P < 0.001, n.s:not significant. (G) UMAPs displaying the expression levels of top down-regulated (Krt8, Krt8 and Csn2,Lmna) and up-regulated (Cdkn2a, Fn1, Lgals1 and Vim) genes in p16+ cycling cells compared to LP and Avd. log10 expression levels are color-coded.

We collected 17,330 high-quality cells in total on which we applied unsupervised graph-based clustering followed by dimension reduction methods to identify cell populations. We conducted a coarse-grained cluster annotation using well-established canonical markers and identified four major cell compartments - *immune* cells, *fibroblasts*, *endothelial* and *epithelial* cells (see Methods, Fig. S1C-F). As our focus was on epithelial cells, we performed a high-resolution sub-clustering on the epithelial compartment (n=11,113 cells, Fig. 1B-C). Top expressed genes per cluster were intersected with a set of known markers of physiological cell populations of the mammary gland (Bach et al. 2021, 2017; Watson and Khaled 2008) (Fig. 1C and Fig. S1F). In the case of clusters composed with cells from either tumor or lesional samples, we labeled them by concatenating the top expressed gene per cluster, with the major sample name which composes the cluster (Fig. 1C and Fig. S1F).

With the objective to map cells undergoing early steps of Brca1-tumorigenesis, we focused on the epithelial sub-compartment prior to tumor detection (Fig. 1C-D, n= 1,706 cells). In 2.7-, 3-, and 5.2- month samples, we identified physiological cell populations of the mammary gland: basal cells (*Krt5*) and clusters of luminal cells (*Krt8*) - luminal hormone-sensing (Luminal H-S, *Prlr*), luminal progenitor (LP, *Aldh1a3*) and secretory alveolar cells (Avd, *Csn2*), with low batch effect within controls (Fig. 1B). The abnormal presence of secretory alveolar cells in the mammary gland of virgin mice at all timepoints (Fig. S1G), confirmed the abnormal differentiation of luminal progenitors into alveolar cells in Brca1/Trp53 deficient mammary glands, which had recently been observed during Brca1 tumorigenesis (Bach et al. 2021).

### Identification of a p16-high cycling population of luminal cells with mesenchymal markers

Apart from these expected cell populations, we identified a cluster of cells, in between normal luminal compartments (LP & Avd) and tumor cells (Fig. 1C-D), characterized by an unequivocal activation of *Cdkn2a*/p16 compared to both LP and Avd (adj. p-value < 2.5e-47, Fig. 1E, Table S1). This partition originated mainly from pre-tumoral glands with lesions (at 5.2 months of age) (Fig. S1G, adj.p value < 0.05, Fisher’s test), but few cells also belonged to mammary glands of 2.7 and 3 month-old animals. As opposed to epithelial cells from the healthy virgin mice individuals (2.7 & 3.2 months), cells from p16+ cluster are cycling just-like tumor cells (Fig. 1F), implying they have by-passed the cell-cycle blockade imposed by p16. In line with p16 activation - a marker of senescence (Collado and Serrano 2010; Koppelstaetter et al. 2008; Di Micco et al. 2021; Campisi and d’Adda di Fagagna 2007) - the transcriptional profile of these cells is significantly enriched for senescence-related hallmark signatures (REACTOME_Senescence Associated Secretory Phenotype, adj. p-value < 2.0 10^-2^). In addition, cells from p16+ cluster express a pro-senescence secreted factor, Igfbp4 (adj. p-value =3.3 10^-23^), that can trigger senescence in neighboring cells (Severino et al. 2013). Our data suggest that these cells, now cycling, may have previously undergone a G1/S blockade and senescence-like state (Buj et al. 2021; Herranz and Gil 2018). Combined over-expression in these cells of *Cdk4* and *Ccnd1* (adj. p-value<2.0 10^-5^), that together promote G1 to S transition, could for example help the cells bypass cycle arrest imposed by p16 overexpression (Roupakia, Markopoulos, and Kolettas 2021).

In terms of identity, these cells show a significant down-regulation of genes characteristic of luminal identity, compared to LP and Avd cells - e.g *Krt8*, *Krt18, Csn2 (Pervolarakis et al. 2020; Bach et al. 2017)* (Fig. 1G). In addition to the repression of epithelial cytokeratins, a series of transcriptional changes testify of dampened epithelial characteristics and acquisition of mesenchymal features: (i) upregulation of *Vim*, *Fn1* and *Sparc* (adj. p-value < 1.5e-17, Table S1), indicative of changes in cytoskeleton and extracellular matrix, and (ii) down-regulation of *Cdh1* and several Claudin genes (*Cldn4*, 3 and 1, adj. p-value < 3.0 10^-8^, Table S1), indicative of the dissolution of adherens and tight junctions. Part of the transcriptional changes could be driven by the transcriptional factor Twist1, that is significantly over-expressed in cells from the p16+ cluster (adj. p-value= 3.0 10^-14^, log2FC=0.45), and is a known activator of *Fn1* and *Sparc*, and repressor of Claudins and E-cadherin genes (Lamouille, Xu, and Derynck 2014), (Fig. 1H, Fig S1H, Table S1). In addition, cells from p16+ cluster display a specific downregulation of *Lmna* (Fig. 1H), indicative of diminished nuclear stiffness, potentially increasing their migration potential (Harada et al. 2014). Altogether, we have identified a population of luminal cells expressing several mesenchymal markers, thereby potentially undergoing a partial EMT (Pastushenko et al. 2018), while aberrantly expressing p16, that we define as a ‘p16 pre-tumoral’ state. The signature of such pre-tumoral state is further kept in tumor cells, as several EMT and senescence-related genes remain over-expressed in tumor cells (see ST sample, Fig. 1G-H and Fig. S1H).

### p16 pre-tumoral state: a transition between luminal progenitor and tumor states

We then sought to reconstitute the timeline of events, and better characterize the transformation from the luminal to tumor state using our single-cell transcriptomics datasets. In order to investigate the presence of a potential continual progression in the epithelial compartment (Fig. 1D), we applied Potential of Heat-diffusion for Affinity-based Trajectory Embedding (PHATE); a non-linear dimension reduction known to efficiently retrieve high-dimensional trajectory structures without specifying any root state (Moon et al. 2019) (Fig. 2A). The non-linear progression between the luminal towards the tumor cells was apparent in the PHATE two-dimensional space (Fig. 2A). p16 pre-tumoral cells were the intermediate between the two pools of cells (normal-like and tumoral), and represented a “bridge” between the two cell populations (Fig. 2A). We further mapped and quantified connections between cell states by performing unsupervised Partition-based Graph Abstraction (PAGA) (Wolf et al. 2019) analysis on the same dataset. According to PAGA representation, the topology of the graph showed that the p16 pre-tumoral node was the most connected, and tumor cell nodes were only reachable through this central hub (Fig. 2B). The highest connectivity score was between the LP and Avd partitions (0.8), corroborating the previously described abnormal LP differentiation to Avd (Bach et al. 2021). The second and third most connected nodes were between the LP and p16 pre-tumoral, and Avd with p16 pre-tumoral partitions respectively, suggesting that the transient p16 pre-tumoral state may arise from either population (Fig. 2B).

**Figure 2:**
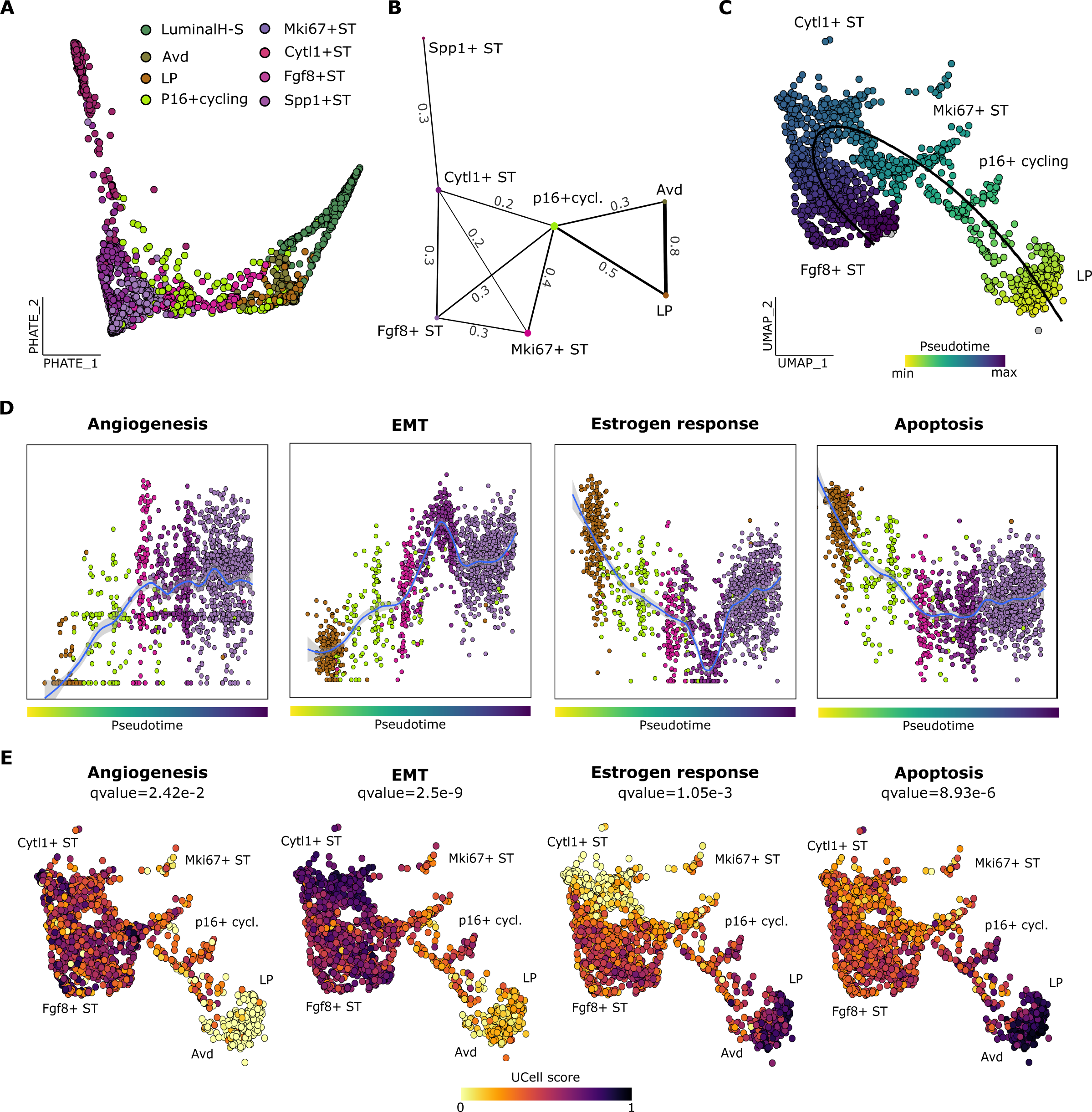
p16-high cycling state is a transitory state between LP and tumor cells. (A) Potential of Heat-diffusion for Affinity-based Trajectory Embedding (PHATE) dimension reduction applied on the zoomed epithelial cells displayed in Fig1C. (B) Partition-based graph abstraction (PAGA) graphical representation of the transitioning clusters, previously represented on UMAP embeddings (Fig1E); nodes are the cell groups and the edge thickness quantifies the connectivity scores between the graph-partitions, highlighted on the graph. (C) UMAP representation with cells colored according to inferred pseudotime values, using the Slingshot algorithm. Transition path is shown by the passing-by line on the cells of interest. (D) Scatter plot representation of transcriptional signatures, based on the gene sets correlated to pseudotime; cells are rankedc by increasing pseudotime values and colored according to their cluster ID. Scores for transcriptional signatures were calculated using UCell (see Material & Methods). (E) UMAP representation with cells colored according to UCell scores.

To decipher the origin of this transient population, semi-supervised pseudotime inference was conducted using the Slingshot algorithm (Street et al. 2018). We hypothesized that either luminal progenitors or alveolar cells could be the starting point, and thus iteratively set them as the root of the tree (Fig. S2A-B). In both scenarios, whatever the root, reaching the ST tumor clusters passes through the p16 pre-tumoral state (Fig. S2A-B). However, when the Avd was set as the root, the algorithm suggested a lineage form Avd to LP, and from Avd to Luminal H-S, which were both biologically irrelevant (Cristea and Polyak 2018; Visvader and Stingl 2014). Therefore, we chose the LP population as the root of the tree, and selected the longest lineage path to reach the Fgf8+ ST cluster (Fig. 2C). Altogether, these complementary approaches model that LP cells switch to a p16 pre-tumoral state prior to tumor formation and growth.

Hereafter, to identify the biological pathways driving this state transition, we studied the top genes correlated to pseudotime values (Fig. S2C; Methods), on which consecutive MsigDB hallmark (Liberzon et al. 2015) pathway enrichment and signature quantification were performed (see Methods). In Fig. 2D-E, we display the most significantly enriched pathways, and transcriptional scores are plotted along pseudotime. While transitioning from LP to tumor state, p16 pre-tumoral cells activate a transcriptional signature associated with angiogenesis and EMT, while inhibiting pathways of apoptosis and estrogen response (Fig. 2D-E). All these four pathway score levels were maintained in all tumor cells (Fig. S2D). Such signatures endorse the pre-tumoral nature of the p16 pre-tumoral state: angiogenesis and inhibition of apoptosis are canonical hallmarks of cancer cells (Hanahan and Weinberg 2016) - meant to enable fast growth of cells, while EMT activation has already been proposed as a mechanism to escape cell cycle arrest *in vitro* (Ansieau et al. 2008; Fridman and Tainsky 2008).

### p16 pre-tumoral signature is specific to basal-like human cancers that rarely recur

We next investigated whether we could detect such transcriptional changes in human breast cancers, and if so which ones. We took advantage of the largest, publicly available, breast pan-cancer bulk RNAseq cohort in TCGA (Berger et al. 2018), to investigate transcriptional similarities between the p16 pre-tumoral state and human breast cancer subtypes. Normalized *CDKN2A*/p16 expression level was the highest in basal-like subtype samples (n=171), as compared to the remaining breast cancer subtypes (Fig. 3A), shown that p16 activation is specific to the basal-like cancers, not only BRCA1-deficient basal-tumors, and a frequent event in this subtype (n=128, 75% from the total basal-like samples). Cross-comparison between the top markers of the basal-like subtype in humans and the top markers of the p16 pre-tumoral cells highlighted the over-expression of EMT-related genes in both populations (*SLPI*, *COL2A1*, *SERPINE2* and *SPP1*) (Fig. S3A). Similarly, *CDKN2A* was the top overexpressed gene in both comparisons, whereas *CSN2* was the top down-regulated gene (Fig. S3A). Altogether, these observations endorse the transcriptional similarity between the p16 pre-tumoral mouse cells and basal-like breast cancers in humans. We further defined a p16 pre-tumoral associated signature, as the top over-expressed genes (log2FC> 0.8 and adj. p-value < 0.05) in the p16 high cluster identified above as compared to both LP and Avd compartments (see Methods). We calculated a p16 pre-tumoral signature score for each sample and observed that basal-like tumors displayed higher scores than other breast cancer subtypes (Fig. 3B); among basal-like tumors, *BRCA1* deficient tumors displayed slightly higher scores than BRCA1 WT tumors.

**Figure 3:**
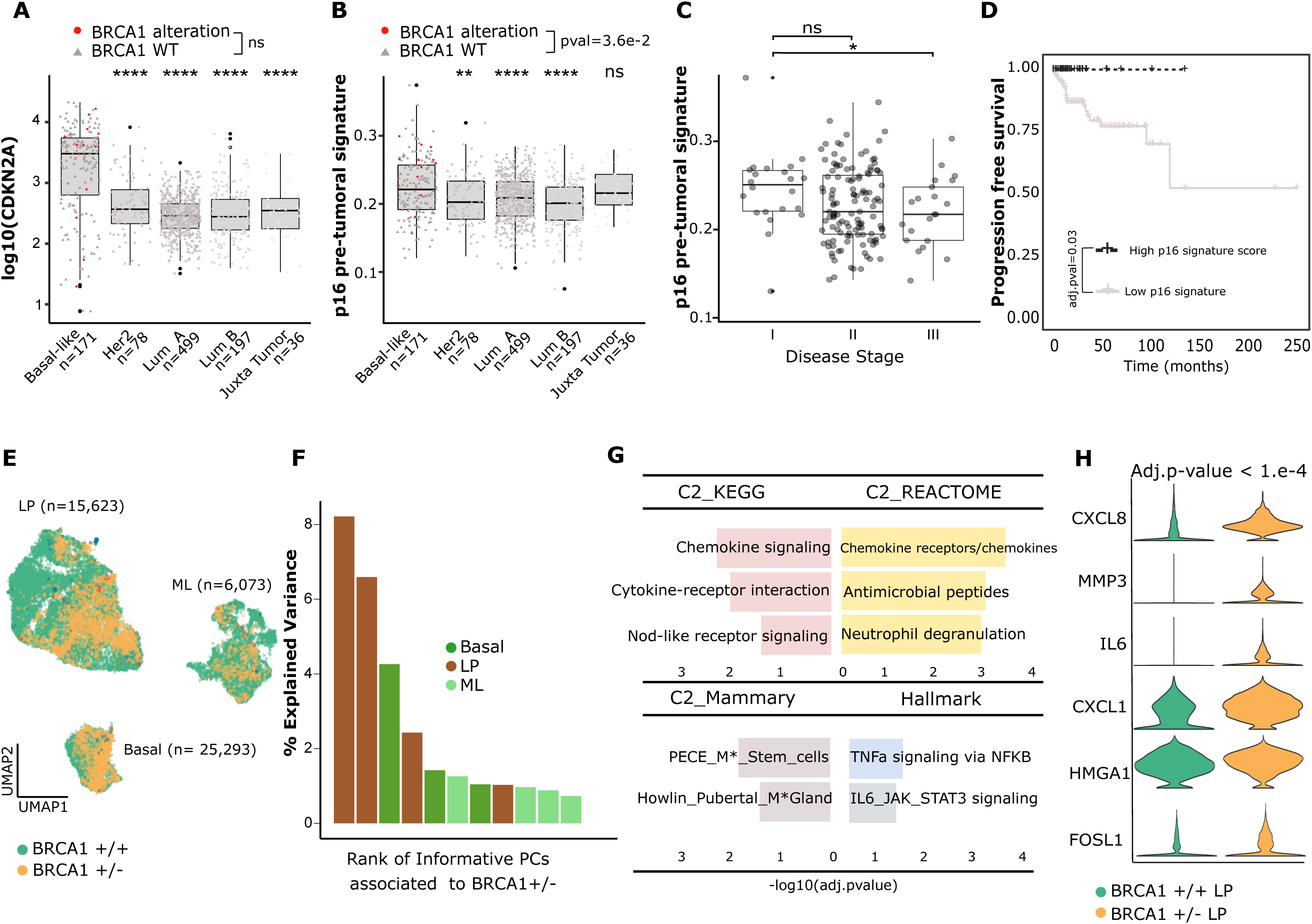
Pre-tumoral signatures in human breast cancers. (A) Boxplot distribution of log10 normalized expression levels of CDKN2A for Breast Pan Cancer TCGA Cohort according to tumor type. (B) Boxplot representation of scores for the p16 pre-tumoral signature according to tumor type. (C) Boxplot representation of scores for the p16 pre-tumoral signature according to tumor stage.(D) Kaplan-Meier disease free survival curve for basal-like tumors, according to score of expression of the p16 pre-tumoral signature. (E) UMAP representation of the epithelial compartment from healthy (N=6) and BRCA1 +/- pre-neoplastic (N=2) human samples, including luminal progenitors (LP), mature luminal (ML) and basal cells; cells were colored according to the sample type of origin. (F) Barplot representation of the top most informative principal components (PC) separating pre-neoplastic from normal-like epithelial cell types. PCs were ranked according to explained variance in each epithelial compartment. (F) Barplot representation of activated pathways (using Hallmark and C2 MsigDB terms) in BRCA1+/- versus BRCA1+/- luminal progenitor compartment, y-axis represent -log10 adjusted p-values. (H) Violin plot representation of expression levels of genes discriminating BRCA1+/- from BRCA1+/+ LP cells.

In addition, p16 pre-tumoral signature was detected in juxta-tumoral samples, and was also more pronounced in early-stage (I) than late-stage tumors (III) (Fig. 3C). Finally, patients who displayed high expression scores of the p16 pre-tumoral signature exhibited longer progression-free survival, (adj. p-value= 0.03), as compared to patients with a low expression score of the p16 pre-tumoral signature (Fig. 3D, S3B-C).

Overall, using public data, we demonstrate that the p16 pre-tumoral signature of initial luminal transformation is detected in basal-like breast cancers and we show it is actually characteristic of early-stage basal-like tumors that rarely recur. These results suggest that in humans, as in mice, this p16-high transcriptional signature corresponds to an early stage in tumorigenesis.

We next investigated pre-tumoral gene signatures prior to tumor formation in humans in a *BRCA1*-/+ deficient context, to understand whether we could detect premises of the p16 pre-tumoral state. To do so, we interrogated gene signatures in the epithelium compartment of patients with germline *BRCA1* heterozygous deficiency (*BRCA1*+/-) prior to tumor formation. We exploited the publicly available scRNAseq human dataset harboring normal-like and *BRCA1+/-* pre-neoplastic mammary gland samples (GSE161529) (Pal et al. 2021). In this context, in contrast to established human tumors and the mouse model presented above, only one copy of *BRCA1* is deficient and *TP53* is initially functional.

We performed the same semi-manual annotation procedure as in the first part and focused solely on the mammary epithelial compartment from normal-like and *BRCA1*+/- pre-neoplastic samples (Fig. 3E, Fig. S3D-E). We projected each epithelial subtype in an independent Principal Component Analysis (PCA) space, and showed that LPs are the cell type most affected by *BRCA1* deficiency (see Methods): the informative PCs with the highest explained variance were retrieved from the LP PCA projection (Fig. 3F, FigS3F-G). LPs in *BRCA1+/*- patients display transcriptional defects, they aberrantly activate genes involved in mammary stem cell signatures and several genes defining a senescence associated secretory phenotype SASP, including chemokines, IL6 and MMP3 (Fig. 3G-H, Table S2). Such analysis confirms, as shown by others, that *BRCA1* deficiency can lead to senescence-like states (Sedic et al. 2015). However, we could not detect, as in juxta-tumoral tissues or human basal-like tumors above, key components of the p16 pre-tumoral state, such as EMT actors or p16 activation. We postulate that such transcriptional changes occur with the onset of tumorigenesis, and probably following *TP53* inactivation (Martins et al. 2012).

### Spatial and temporal analysis of the p16 pre-tumoral state

We next sought to validate and spatially resolve the acquisition of the p16 pre-tumoral state in the mammary gland in *situ*. We performed multiplex immunohistochemistry on paraffin-embedded formalin-fixed (FFPE) sections from mice at different stages of tumorigenesis, with either normal-like epithelium, lesions, or tumors (including both Cre- and Cre+ animals) (Fig. 4A). This technique aimed to simultaneously monitor, on each section and at the single cell resolution, for over 70,790 cells: i) cell identity (Krt8, Krt5), ii) cell cycle status (p16, Ki67), and iii) epithelial to mesenchymal plasticity (EMP) (E-cadherin, N-cadherin, Vim) in addition to hematoxylin staining (Fig 4A, Fig S4, see Methods). We first quantified the proportion of p16 positive (p16+) cells within the epithelial compartment across timepoints starting at 3 months. We detected p16+ cells (>5% in average) within lesions and tumors (Fig. 4A-B), confirming our single-cell transcriptomic analyses (Fig. 1). Importantly, while we did not observe any p16+ cells in the Cre- animals (Fig. S4C), they were found within normal-like Cre+ ducts as early as 3 months (upper panel Fig. 4A), as well as within juxta-tumoral tissues in tumor-bearing animals. At the earliest time point, prior to any lesion or tumor formation, the p16+ cells are isolated single cells (Fig. 4A, S4B) and located in the inner part of the duct, within the luminal compartment. Such initial localization, in addition to the correlation of p16 and Krt8 staining (Fig. 4C) confirmed their luminal origin (89% of the total p16+ cells).

**Figure 4:**
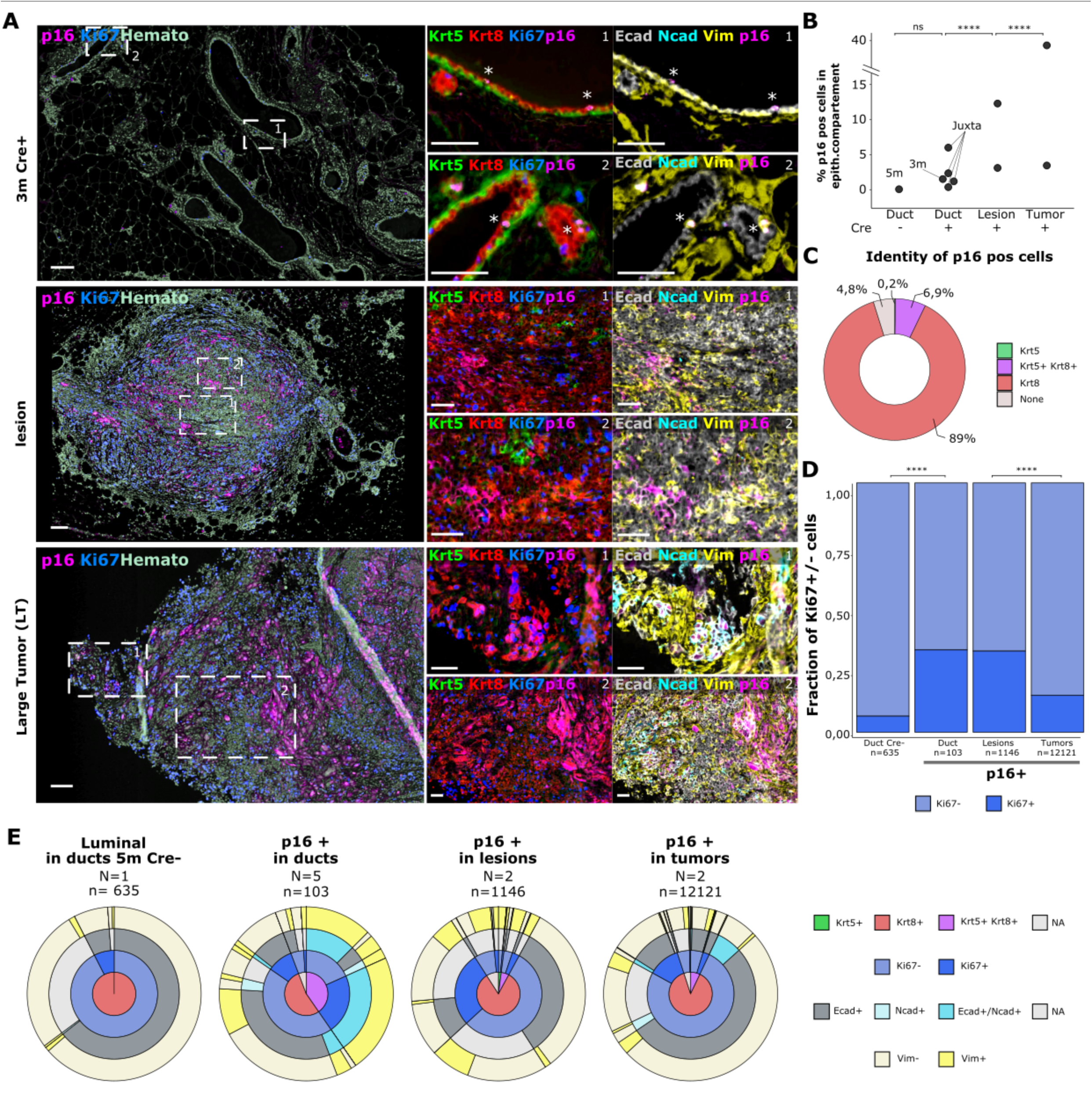
Spatial characterisation of p16 activation and partial EMT. (A) Left: Representative IHC staining for p16 (pink), Ki67 (blue) in young normal-like mammary epithelium from Cre+ 3-month-old mouse, in gland with lesion and in tumor, scale bar represents l00µm. For zooms, sections are stained for identity markers Krt5 (green), Krt8 (red), cell cycle markers Ki67 (blue), p16 (magenta) and EMT markers Ecad (white), Nead (cyan), Vim (yellow), scale bar represents S0µm. (B) Dot plot representation of the percentage of p16+ positive cells at different stages of mammary tumorigenesis, including a negative control “duct Sm Cre-ℍ and normal-like ducts from lesion-free or juxta-tumoral regions (N=S Cre+ in total), N=2 samples with lesions, and N=2 tumor samples. Asteriks above each sample category represent the significance levels of pairwise comparisons between the 4 categories (Cre+ vs Cre-, lesional vs duct Cre+ and tumor vs lesional samples, respectively) with Fisher’s test, *p < 0.05, **p < 0.01, ***p < 0.001. (C) Donut plot representation of the epithelial subtype origins of the pl6+ cells, epithelial classes were color-coded. (D) Barplot displaying the fraction of Ki67+/- cells within p16+ cells fractions, and within a control population of luminal cells from Sm Cre-mouse. (E) Sunburst plot representation of the hierarchical distribution of control luminal cells from Sm Cre-mouse and p16+ cells, according to their mammary ID, cell-cycle status, and expression of EMT markers.

We next wanted to assess the proliferative capacity of these p16+ cells using Ki67 staining. In both duct and lesional samples, more than 32% of p16+ cells are Ki67+, supporting their capacity to escape p16-mediated cell cycle arrest at very early time-points. The proliferative index of p16+ cells is significantly higher in duct and lesions, compared to p16+ cells in tumors (15% Ki67+) and control mammary epithelial cells (6% Ki67+), suggesting that p16+ cells at the onset of Brca1 tumorigenesis are particularly proliferative (Fig. 4D).

Finally, we interrogated the extent of EMP within the epithelial compartment using Ecad, Ncad and vimentin stainings (Fig. 4E-F). In control Cre- mice at 5 months, luminal cells mostly display expected E-cadherin junctions and no vimentin-based filaments (Fig. 4F left donut). In contrast, p16+ cells within the luminal compartment of Cre+ animals were significantly enriched for vimentin and displayed enrichment for N-cadherin - together (40%) or not with E-cadherin (48%) (Fig.4E 2nd donut). In addition, they frequently displayed both Krt8 and Krt5 staining. Dual enrichments for E-cadherin and N-cadherin were not observed in lesions but in rare cases in tumors, underlying the metastable nature of EMP. In tumors, p16+ cells were grouped into spans of neighboring cells with high N-cadherin and vimentin expression (Fig. 4A, bottom panel, Large Tumor), confirming our observations by scRNAseq that only a fraction of tumor cells remain p16+ and harbor strong expression of mesenchymal features (Fig. 1G-H).

Complementary to our initial single cell transcriptomics analyses, multiplex imaging enabled the extensive search for p16 rare pre-tumoral states in whole tissue slides at various time points. It allowed to detect p16+ individual isolated cells within the inner part of the mammary duct with mesenchymal markers at early time points, before any lesion or tumor formation, suggesting EMP is an early event in basal-like tumorigenesis. Losing epithelial characteristics could be essential for the rupture of the duct structure and formation of the initial tumor bud.

### p16 pre-tumoral state occurs post genomic crisis

Loss of BRCA1 impairs homologous repair (HR) mechanisms, and leads to a major genomic crisis, with the accumulation of multiple genomic alterations (Scully and Livingston 2000; Polak et al. 2017). In such a context, we thought it was critical to understand when such a genomic crisis was occurring, and how it related to the phenotypic switches we were observing. We used our scRNAseq epithelial dataset to quantify genomic rearrangements and to investigate clonal evolution across cells and time points. Copy Number Variation (CNV) were first inferred from the scRNAseq data using inferCNV (Patel et al. 2014), taking the basal cells as reference, as the Cre is not expressed in these cells (Molyneux et al. 2010). For each cell, we calculated the percentage of their genome displaying genetic alterations (Fig. 5A-B, Fig. S5A). All tumor cells display similar percentages of rearranged genomes (Fig. 5B, median= 35.4%), whatever their size, suggesting that the genomic crisis probably occurred prior to tumor expansion. LP cells of tumor-free & lesion-free animals already display a high percentage of CNVs (median 23 %) compared to the basal cells (median 5.3%), even at 2.7 months (median 22.4%). Such observations imply that the LP compartment can tolerate numerous CNVs following Brca1/Trp53 deletion, without any rapid phenotypic consequence. Rates of genome rearrangement in p16 pre-tumoral cells are among the highest of the LP compartment (median % alteration = 25%), yet their maximum rate does not exceed what is observed in the LP population (Fig. 5B). Altogether, we show that the major outburst of CNVs occurs in the LP compartment prior to any tumor formation, in agreement with previous studies which showed that copy number alterations were acquired in short punctuated bursts at early stages of tumor formation (Gao et al. 2016). These results suggest that the genomic crisis triggered by Brca1/Trp53 deletion is not sufficient to launch tumorigenesis, and precedes the partial EMT processes identified above.

**Figure 5:**
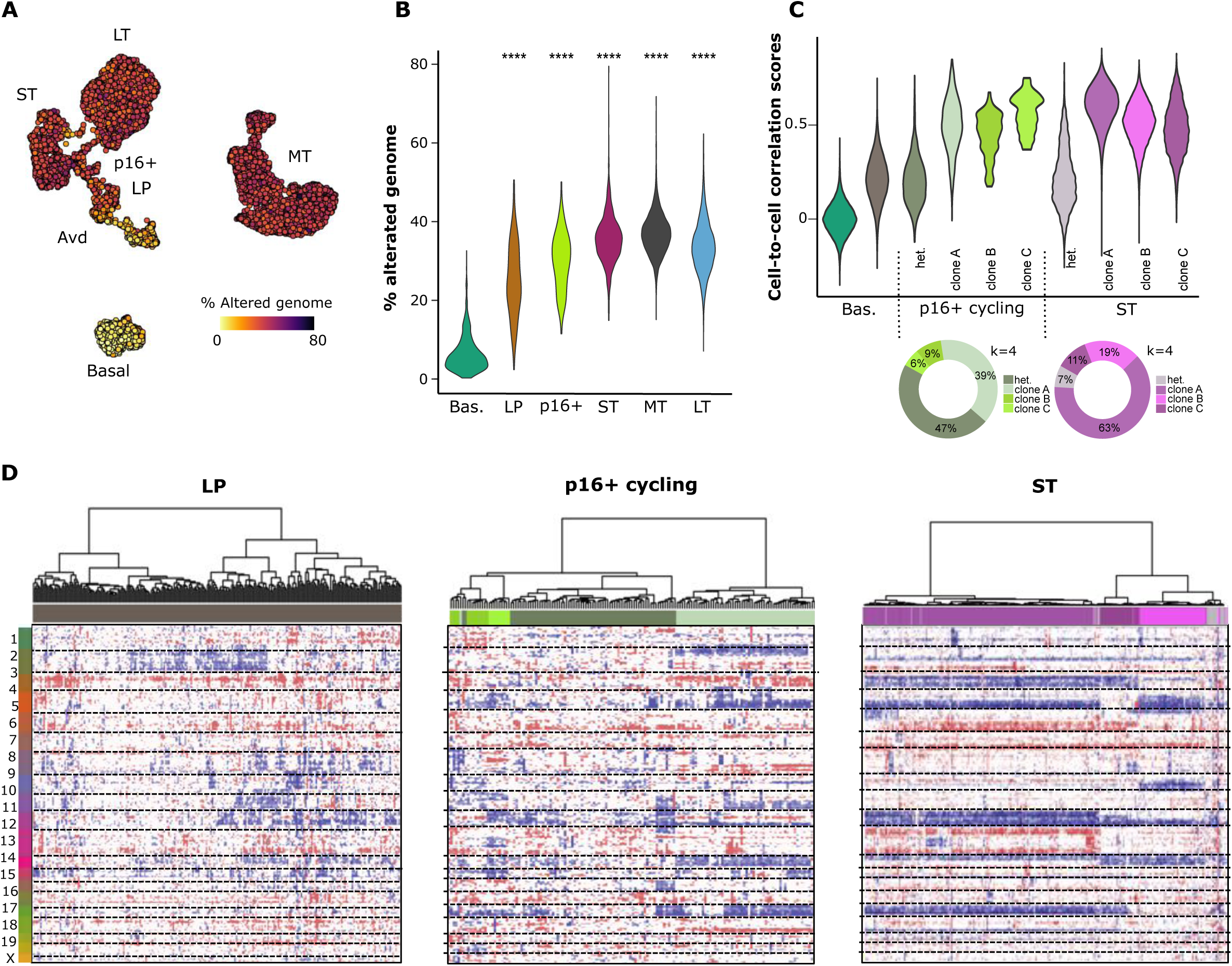
Clonal evolution from LP, to p16-high cycling populations and tumors. (A) UMAP representation of the transcriptomes of cells from the epithelial compartment with cells colored according to the percentage of their genome with CNV. (B) Violin plot distribution of the percentage of altered genome per cell, grouped by cluster ID; horizontal lines represent the median values. Asterisks represent the significance levels of mean comparison with basal cells. *P < 0.05, **P < 0.01, ***P < 0.001. (C) Upper panel: violin plot distribution of the pairwise intra-cluster correlation scores between single-cell CNV profiles. Cluster partition (i.e clone separation) was achieved with Consensus Clustering (Fig. S3). For LP and Basal clusters, intra-cluster correlation scores were computed on all cells of each compartment due to the absence of any consensual optimal number of clusters; Lower panel: Donut plot representation of the number of cells per clone for p16+ cycling cells and ST cells. (D) Heatmap representation of log-transformed residuals from inferCNV, with basal cells as a reference, for LP, p16+ cycling cells and cells from ST sample; blue and red values refer to deletions and gains respectively. Horizontal dotted lines separate chromosomes.

### p16 pre-tumoral state is multi-clonal

To understand how p16 pre-tumoral cells emerged from the LP compartment - through clonal expansion or a few cells or state transitions in multiple LP cells - we next investigated clonal dynamics within the LP compartment, p16 pre-tumoral population and the small tumor. We performed integrative consensus hierarchical clustering to identify genetic clones within each cell population (see Methods); and samples with no stable partition were considered as highly multi-clonal (Fig. S5B). In addition, we evaluated correlation scores between single-cell CNV profiles across clusters, to further confirm the absence or existence of sub-clones; considering that genetic clones will show high intra-correlations scores by definition.

As expected, we could not partition the population of reference basal cells into clones, further confirmed by a random distribution of cell-to-cell correlation scores (Fig. 5C). In the LP compartment, we could not identify any stable partition (Fig. 5D and Fig. S5B-C) into clones, further supported by low cell-to-cell correlation scores (Fig. 5C). In the p16 pre-tumoral cells, 47% of cells remained highly multi-clonal, with cell-to-cell correlation scores similar to those of the LP compartment, but we could also identify 3 clones accounting for 53% of cells (Fig. 5C-D, Fig. S5B,D). In contrast, the small tumor was organized into 3 major clones (Fig. 5C and Fig. S5E) accounting for 93% of the cells (Clones 1, 2 & 3), with high intra-clone correlations scores (median 0.56) (Fig. 5C-D). These results suggest that the transition from the LP to p16 pre-tumoral state can be achieved by a multitude of cells, and not only by isolated clones that are being selected for. This strongly suggests the contribution of non-genetic mechanisms to this transition state, potentially dedifferentiation or partial EMT mechanisms, identified above.

### Disruption of heterochromatin at the onset of tumorigenesis

To further characterize the p16 pre-tumoral state, we next investigated canonical markers of senescence associated to p16 upregulation (Collado and Serrano 2010): presence of B-galactosidase (Bgal) and senescence-associated heterochromatin foci (SAHF) within tissues. We could not quantify any Bgal within juxta-tumoral or tumor sections (Fig. S6A), however we identified SAHF-like structure in lesions by immunofluorescence (Fig. 6A). Regarding SAHF, they were initially defined as main cores enriched in H3K9me3 mark, coated by enriched rings in H3K27me3 (Aird and Zhang 2013; Paluvai, Di Giorgio, and Brancolini 2020). As H3K9me3 mark is already organized in foci in non-senescence cells in mice -chromocenters (Probst and Almouzni 2008), we chose H3K27me3 staining to study changes in heterochromatin organization during Brca1 tumorigenesis.

**Figure 6:**
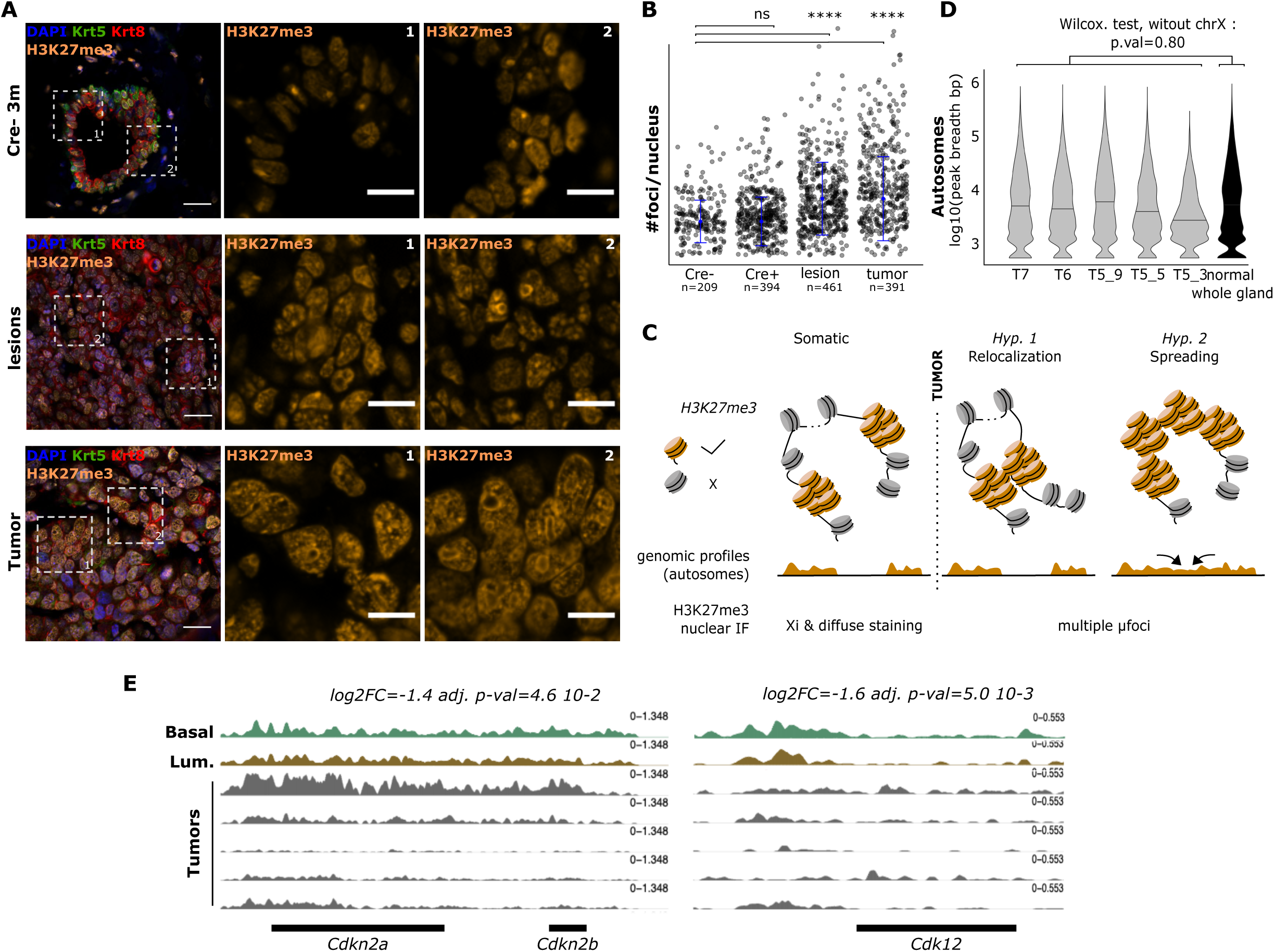
Heterochromatin disruption. (A) Representative sections for mammary gland from 3 months-old Cre-mouse, 5 months old Cre+ mouse with lesion, and tumor. All are stained by immunofluorescence for basal marker Krt5 (in green), luminal marker Krt8 (in red), histone mark H3K27me3 (in orange), Dapi (in blue) (left image in each panel), scale bars represent 20μm. For zooms scale bars represent 10μm. (B) Jitter plot representation of the number of μFoci per nucleus in the studied samples in (A). Asterisks represent the significance levels of median comparison with the Cre -/- control sample. ns: non-significant, *P < 0.05, **P < 0.01, ***P < 0.001. (C) Scheme showing the two hypotheses that could explain the observations of H3K27me3 μfoci. (D) Violin plot of the H3K27me3 peak breadth on autosomes in tumor samples compared to normal-like mammary glands. (E) Cumulative coverage plot for H3K27me3 signal in Cdkn2a/b,Cdk12,promoter genes in sorted basal and luminal population and tumor samples.

As expected, in mammary gland controls from Cre-mice, H3K27me3 staining revealed one single foci per cell (Fig.6A-B), corresponding to the inactive X (Xi), whereas the remaining staining is homogeneously diffused in the nucleus (Fig. 6A, Fig. S6B top panel). In lesions and tumors, we observed the accumulation of multiple H3K27me3-enriched chromatin forming small aggregates, that we termed micro-heterochromatin-foci (µ-HF) (Fig. 6A, Fig. S6B). In addition, H3K27me3-enriched chromatin tended to accumulate in ring-like structures, surrounding nuclear regions devoid of DNA as attested by negative DAPI staining-possibly corresponding to nucleoli (Cmarko et al. 2008).

We further investigated the special case of the heterochromatin of the Xi, as the loss of the Barr Body is a hallmark of basal-like breast cancers, whether through a genetic loss of the inactive X chromosome (Ganesan et al. 2004; Vincent-Salomon et al. 2007) or following epigenomic reprogramming and reactivation of the inactive X (Chaligné et al. 2015). Using both H3K4me3 and H3K27me3 tumor derived datasets, we actually show that H3K27me3 signal from the X-chromosome is lost in tumor cells, indicating that the Xi is either genetically lost or it has lost its repressive chromatin enrichment, in both cases attesting disruption of this heterochromatin structure (Fig. S6D). The absence of H3K4me3 enrichment anywhere on the X chromosome in these same cells, demonstrates the absence of partial or total reactivation of the Xi, and favors the genetic loss of the Barr Body in the cells (Fig. S6D-E).

We hypothesized (Fig. 6C) that the accumulation of µ-HF could either be the result of (i) the nuclear reorganization of regions of heterochromatin, leading to co-localization of multiple heterochromatic regions, or (ii) the expansion of H3K27me3 enrichment on large genomic regions - similar to Xi-enrichment. We used H3K27me3 genome-wide maps to test both hypotheses as only the latter would lead to genomic redistribution of H3K27me3 marks. We generated H3K27me3 ChIPseq datasets for n=5 tumors and compared them to published datasets for normal mammary cells (Pal et al. 2013). When comparing H3K27me3 peak breadth genome-wide, tumors did not show largest H3K27me3 peaks on autosomes (Fig. 6D). These results demonstrate the absence of large heterochromatinization phenomena on autosomes (>Mb), and suggest that formation of µ-HF rather corresponds to spatial reorganization of existing chromatin regions.

We next investigated whether this major nuclear reorganization was associated with focal H3K27me3 changes during tumorigenesis, undetectable at the microscopic scale but at the genomic scale. We included in our analysis H3K27me3 profiles of published FACS-sorted mammary gland luminal and basal cell populations, along with our tumor samples to seek for recurrent epigenomic differences between normal and malignant samples. Principal component analysis (PCA) showed that tumors have heterogeneous repressive epigenomes, yet 34% of variance is driven by common tumor-specific epigenomic features (PC1) (Fig. S6C). When comparing tumors versus cells of the physiological gland, we show that several cell cycle genes (*Cdkn2a*, *Cdk12*, *Cdk6*) display a recurrent loss of repressive H3K27me3 enrichment in tumors (Fig. 6E), both inhibitors and promoters, suggesting that local epigenomic remodeling could participate both in the entry and exit of cell cycle during tumorigenesis. Loss of H3K27me3 had already been shown to enable *Cdkn2a* transcriptional activation in early senescence (Ito et al. 2018).

Altogether we show that disruption of heterochromatin - with a drastic spatial reorganization in the nuclei and rare local rearrangements - occurs early in tumorigenesis, potentially as a consequence of a senescent-like state.

## DISCUSSION

Our study provides a detailed mapping of the transcriptional, genetic and epigenetic evolution of epithelial cells during early stages of basal-like breast tumorigenesis *in vivo*. Thanks to a mouse model launching tumorigenesis in multiple luminal progenitor cells, we have been able to detect rare state transitions occurring in epithelial cells prior to tumor formation - that cannot be studied in humans. Our *in vivo* results partially bridge the gap between observations from pre-tumoral tissues and established basal-like tumors in humans (Figure 7). We show the occurrence of epithelial to mesenchymal plasticity (EMP) in the luminal compartment of mammary glands at the onset of tumorigenesis. Our data demonstrate that luminal progenitor cells can switch to a p16+ cycling state, with an activation of partial EMT and angiogenesis-related pathways while shutting down apoptosis and estrogen-related signaling. We propose that these cells have previously undergone a transient cell cycle arrest, supported by two features of the senescent state: (i) *Cdkn2a/*p16 demethylation and subsequent expression and (ii) the drastic reorganization of heterochromatin with the formation of multiple heterochromatin foci. It has previously been shown that loss of Brca1 is followed by senescence-like processes, whether in mammary epithelial cells or even in embryos (Cao et al. 2003; Sedic et al. 2015). It is often triggered by teliomerism or after activation of oncogenes expression and mediated by Trp53. Senescence has also been frequently shown to occur in breast cancer cells following irreversible damage or cancer treatments, preventing the cells from proliferating and thereby stopping the tumor growth (Ewald et al. 2010; Fitsiou, Soto-Gamez, and Demaria 2021). Yet little was known about how and whether these tumor cells escaped such senescence-like phenomena. EMP occurs both in normal and pathological contexts, e.g embryonic development, wound healing, fibrosis, or cancer metastasis, where it enables cells to adopt a migratory and invasive behavior (Nieto et al. 2016). In the mammary gland, it is involved both in organogenesis and cancer metastasis (Chakrabarti et al. 2012). During tumor progression, EMT is known to participate in cancer dissemination, as it enables cells of the primary tumor to leave the tissue of origin through partial dissociation of the primary carcinoma. Our results are one of the first examples *in vivo* of the occurrence of partial EMT at the onset of tumorigenesis. The transition from an epithelial to mesenchymal state is often incomplete and metastable (Pastushenko et al. 2018), with cells in intermediate states combining epithelial and mesenchymal features as we do observe here. Combining single-cell transcriptomics and multiplex imaging in tissues, we have identified several characteristics of EMP (Yang et al. 2020) in luminal progenitor cells as they leave the luminal compartment to form lesions: (i) remodeling of the cytoskeleton, with a decrease in cytokeratins (Krt8, Krt19 and Krt18) and a switch to vimentin-based filaments, (ii) reduced cell-cell adhesive properties with the decrease in E-cadherin expression and apparition of N-cadherin, (iii) the expression of the transcription factor Twist1, and (iv) modifications of the extracellular matrix with the expression of Fibronectin. For the latter, we have also observed a specific expression of type XI Collagen from pre-tumoral cells (*Col11a1*, *Col2a1* a.k.a *Col11a3*). Type XI collagen is characteristic of deregulated matrisome of the most aggressive tumors across cancer types (Nallanthighal, Heiserman, and Cheon 2021; S. B. Lim et al. 2017; Pearce et al. 2018) - whether expressed by tumor or stromal cells, high *COL11A1* is associated with cancer invasiveness and metastasis.

**Figure 7:**
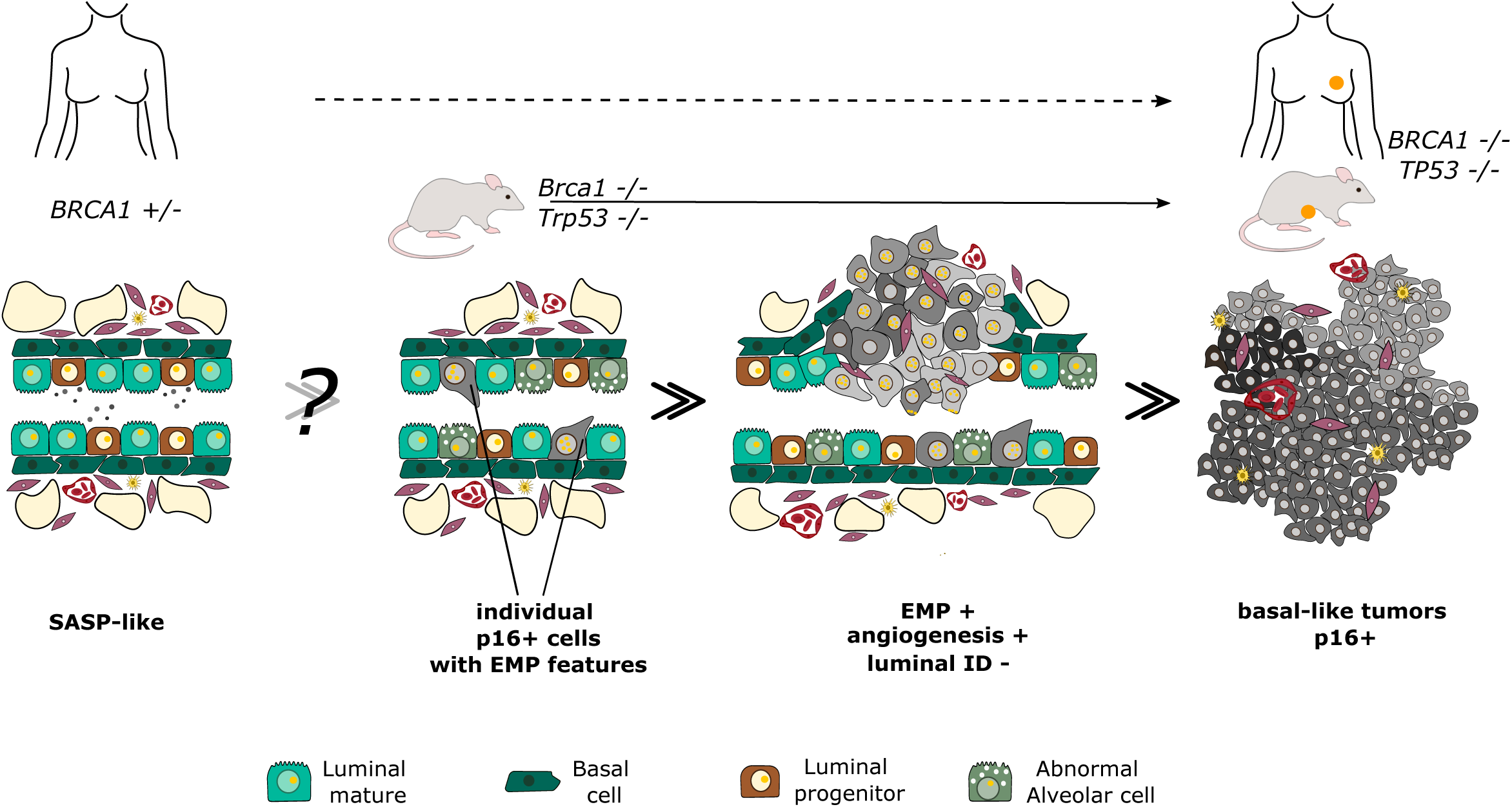
Understanding early steps of BRCA1 tumorigenesis in mouse and human.

What exactly launches EMP in early tumorigenesis remains to be determined. Here, we show that the transcription factor Twist1 is transcribed in the pre-tumoral population; it could orchestrate part of the EMP phenotype that we observe. We have indeed found in pre-tumoral cells signs of Twist1 activity with the activation of its target genes, Fibronectin, N-Cadherin and Sparc, and repression of its known epithelial target genes E-cadherin and Claudin genes (Cldn4, 3 and 1) (Lamouille, Xu, and Derynck 2014). Yet it remains to be determined what exactly triggers its activation. *BRCA1* itself has recently been identified as a guardian of the epithelial states (Zhang et al. 2022) - inactivation of *BRCA1* by CRISPR leads to increased EMP in mammary cells. Another trigger of EMT could also be senescence - itself induced by extensive genomic rearrangements following Trp53 and Brca1 deletion. It has been proposed *in vitro* that EMT, driven by Twist1 and 2, could help override Ras-induced senescence in mouse fibroblasts (Ansieau et al. 2008). Recently, it was shown that senescence actually bridges RAS activation and EMT over the course of malignant transformation in human mammary epithelial cells (De Blander et al. under consideration). In a therapy-induced senescence phenotype, it was also shown that senescence promotes reprogramming and cancer stemness (Milanovic et al. 2018), suggesting that non-genetic mechanisms could be tightly associated to the entry and exit of the senescent state in various contexts. Here, based on our *in vivo* results, we propose that during early basal-like breast tumorigenesis, luminal progenitor cells undergo a cell-cycle arrest characterized by p16 activation, and that subsequently cells bypass this arrest through non-genetic mechanisms, potentially with partial EMT, as demonstrated by the existence of a p16+ cycling population harboring both epithelial and mesenchymal markers.

In the earliest step of tumorigenesis, we have also observed the occurrence of a major heterochromatin crisis, with the total reorganization of H3K27me3-enriched chromatin in the nucleus *in vivo*: a loss of the inactive X (Xi) together with the accumulation of H3K27 foci. The foci resemble senescence-associated heterochromatin foci (SAHFs), hallmarks cellular senescence (Stone, McCabe, and Ashworth 2003; Kristiansen et al. 2005; Sirchia et al. 2005). The destabilization and loss of the Xi could be a consequence of a global 3D reorganization of H3K27me3-enriched heterochromatin in the nucleus. Recruitment of heterochromatin to nucleoli structures - as observed in pre-tumoral and tumoral cells - could for example lead to the destabilization of the Xi (Bizhanova and Kaufman 2021). Such observations - together with the demethylation and subsequent activation of p16 - suggest that these pre-tumoral cells might have entered at some point a senescence-like state. Such epigenomic abnormalities are recurrently observed in full grown basal-like tumors, notably the loss of the Xi through genetic or epigenetic events (Ganesan et al. 2004; Chaligné et al. 2015; Vincent-Salomon et al. 2007). We propose that heterochromatin abnormalities, together with p16 upregulation, in human tumors might be scars of such initial heterochromatin crisis and associated senescence-like state.

Finally, our work opens up several translational perspectives for the early interception of *BRCA1* tumorigenesis and potential patient stratification. Detecting single tumor-initiating events in humans is close to impossible, and we have used a mouse model as a magnifying glass to detail early state transitions in Brca1/Trp53 deficient epithelium. As mentioned in the introduction, in *BRCA1*-germline carriers, *TP53* mutation was actually shown to be among the earliest events in tumor formation (Martins et al. 2012). Using our mouse datasets, we were able to define a p16 pre-tumoral signature, characteristic of the epithelial changes occurring at the onset of basal-like tumorigenesis and kept on during tumor formation. In human tumors, we show that this signature is specific to basal-like cancers, just like p16 overexpression. In addition, we show that it has prognostic potential: with basal-like tumors, patients with high pre-tumoral signature score have a significantly longer disease-free survival. Our results suggest that basal-like tumors with a high pre-tumoral signature score might have been at an earlier stage, hence with a better outcome. Our pre-tumoral gene signatures could constitute candidate biomarkers to detect early epithelial transformation and be favorable prognostic markers. Among the earliest events to detect, we show the advent of dual expression of basal and luminal markers - supported by multiplex imaging data of isolated p16+ cells detaching from the luminal compartment. A recent study actually shows that alveolar cells with dual basal/luminal markers, and a gene-signature associated with basal-like cancers, accumulate with age in human breast (Gray et al. 2022), further highlighting the potential interest of cells with poor lineage definition.

In terms of therapeutic targets, preventing the early state transitions occurring in the luminal progenitor compartment switch from luminal to a pre-tumoral p16+ cycling state for example, could be a relevant therapeutic avenue that we need to investigate. One strategy would be to target EMP, by hampering the listed characteristics above to destabilize this plastic state, with for example COL11A1 inhibitors (Nallanthighal, Heiserman, and Cheon 2021), or by launching Twist1 degradation with harmine (Yochum et al. 2017). Another Achilles heel of the pre-tumoral state could be the mechanisms used to re-entry cell cyclepost cell cycle arrest upon p16 activation. We have shown that pre-tumoral cells over-express both Cdk4 and Ccnd1, that together promote the switch from G1 to S phase, and are antagonized by p16. Such combined over-expression might be a mechanism for these cells to escape p16 overexpression. In this line, p16+ cycling pre-tumoral cells might be particularly sensitive to CDK4/6 inhibitors.

## STAR Methods

### Animal models

he generation of Brca1^fl/fl^ and Trp53^fl/fl^ mice has been previously described (Jonkers et al. 2001; Liu et al. 2007). Blg-Cre transgenic mice were purchased from The Jackson Laboratory. Mice strains were crossed to obtain Blg-Cre Trp53^fl/fl^ Brca1^fl/fl^ animals. Genotypes were determined by PCR (primers Cre: 3’ CGAGTGATGAGGTTCGCAAG 5’ - 3’ TGAGTGAACGAACCTGGTCG 5’; primer Brca1 : 3’TATCACCACTGAATCTCTACC 5’ - 3’ GACCTCAAACTCTGAGATCCAC 5’; Trp53: 3’ AAGGGGTATGAGGGACAAGG 5’ - 3’ GAAGACAGAAAAGGGGAGGG 5’). Mice were sacrificed by cervical dislocation. For each sample (gland or tumor), one piece was fixed in 4% paraformaldehyde (15710, Euromedex) for histological analysis, one piece was snap frozen in dry ice and stored at -80°C and one piece was kept fresh for the desired experimentation.

### Ethics statement

All procedures used in the animal experimentations are in accordance with the European Community Directive (2010/63/EU) for the protection of vertebrate animals. The project has been approved by the ethics committee n°02265.02. We followed the international recommendations on containment, replacement and reduction proposed by the Guide for the Care and Use of Laboratory Animals (NRC 2011). We used as few animals as possible and minimized their suffering, no painful procedures were performed. The breeding, care and maintenance of the animals were performed by the Institut Curie animal facility (facility license #C75-05-18).

### Immunostaining

Glands and tumors were fixed in 4%PFA/PBS at 4°C overnight, then washed with PBS (Gibco, 10010023) a first time for 1h and a second time at 4°C overnight. The samples were then passed through consecutive (50%, 60%, 70%) ethanol baths for 30 min each at room temperature. Paraffin embedding and sectioning (5µm) was performed by the experimental pathology department of Institut Curie. At the staining time, the slides are dewaxed by heating at 65°C for 1h and wash 2 times in Xylene 10min, then rehydrated via consecutive bath: 2x Ethanol 100% (VWR 20821,31) 10min, 1x Ethanol 90% 5min, 1x Ethanol 80% 5min, 1x Ethanol 70% 5min, 1x Ethanol 50% 5min, 2x Water 5min. Retrieval treatment was performed by incubation in citrate buffer (C9999) for 20min at 95°C. After a 1h room temperature cooling, the slides are cleaned in PBS and permeabilized in permeabilization buffer (BSA 2%, FBS 5%, Triton 0,3% in PBS) for 2h at room temperature. Primary Antibody incubation was done on blocking buffer (BSA 2%, FBS 5%, PBS) at 4°C overnight with Chicken Krt5 antibody 1:500 (905901), Rat Krt8 antibody 1:500 (MABT329), Rabbit H3K27me3 antibody 1:20 (C36B11), Rabbit p16 antibody 1:100 (Abcam, ab211542). After 3 washes in PBS for 10 min each, incubation of the antibodies was performed for 2h at room temperature with: goat anti-rabbit Cy3 1:1000 (A10520), goat anti-rat Cy5 1:1000 (A10525), goat anti-chicken Alexa Fluor 488 (A11039) 1:500, DAPI 0,5µg/ml. After 3 wash in PBS 10min, sections were mounted in Aquapoly mount media.

### LacZ staining

Glands and tumors were directly fixed in PFA 4% for 2h and incubated in PBS, 30% Sucrose at least 24h. Samples were included in optimal cutting temperature OCT medium (23-730-751) in moulds and cooled on a metal support previously cooled on dry ice. The samples were stored at - 80°C before being cut in a cryostat at -20°C in a 6µm section. Slides were stored at -80°C before use. For the staining, the slides were equilibrated at room temperature for 10-20 min and washed 3 times for 5 min at room temperature in the washing buffer: PBS, 2mM MgCl2, 1x Na-DOC, 0,02% NP40. After that, slides were incubated in the LacZ Stain: Washing solution, 10mM K3Fe, 10mM K4FE, 1,5 mg/ml X-Gal in a humidified chamber in the dark at 37°C for 4h to overnight. Slides were washed in a consecutive bath of: PBS for 1 min then for 15 min at room temperature, water for 15 min at room temperature and (optionally) Nuclear fast red for 5 min and 2 times in water for 5 min each. Sections were mounted in Aquapoly mount media.

### Microscopy, image acquisition and analysis

Image acquisition of stained sections were done using a laser scanning confocal microscope (LSM780, Carl Zeiss) with a LD LCI PLAN-APO x40 or x65/08 NA oil objective. The acquisition parameters were: zoom 0.6; pixel size xy 554 nm; spectral emission filters (bandwidth): 414-485 nm, 490-508 nm, 588-615 nm, 641-735 nm; laser wavelengths: 405, 488, 561 and 633 nm. Images were captured using Metamorph. Image processing was performed using Fiji Software, version 1.0. The counting of µ-HF was done in Fiji with a custom macro, for each nucleus, we selected the most representative Z, then the counting was done automatically with the AutoThreshold MaxEntropy.

### Multiplex histological staining

Multiplexed IHC was performed according to the protocol developed by (Remark et al. 2016), with some adjustment. Tissues were baked at 60°C for 1h, deparaffinized in Xylene (Fisher Scientific, 10467270) and rehydrated. The heat-induced epitope retrieval was done with pH6.1 citrate buffer (Dako, S169984-2) or pH9 EDTA buffer (Dako, S236784-2) in a 95°C water bath for 30 minutes for the first staining (otherwise 15min) followed by incubation in REAL peroxydase blocking solution (Agilent Dako, S202386-2) for 10 minutes. If the primary antibody was the same species as any antibody used in prior stains, another blocking step was added with Fab Fragment, only for anti-rabbit (Jackson ImmunoResearch Europe Ltd, 711-007-003) during 20 minutes. Protein block serum free (Agilent Dako, X090930-2) was added for 10 minutes. Primary antibody was incubated for 1 or 2 hours at room temperature or overnight at 4°C. The primary antibody was detected using a secondary antibody directed against the first one, conjugated with horseradish peroxydase (Anti-rabbit: Agilent Dako, K400311-2) (Anti-rat: BioTechne, VC005-050) followed by chromogenic revelation with 3-amino-9-ethylcarabazole (AEC) (Agilent Dako, K3468). Slides were counterstained with hematoxylin (Thermo Scientific, 6765001) and mounted with Glycergel aqueous mounting medium (Dako, C056330-2). After scanning (Philips Ultra Fast Scanner 1.6 RA), tissues were bleached with ethanol baths and another cycle was performed starting with the heat induced epitope retrieval.

### Overlay of multiplex histological stainings

Histological analysis was performed using the open-source image analysis QuPath software (QuPath-0.3.2, http://qupath.github.io/) (Bankhead et al., n.d.) and ImageJ/Fiji. We created a new QuPath project containing all scans of each slide which allow us to crop and export (BioFormats plugin) and then overlay the images using Fiji script following these different steps: 1. Color deconvolution (separation of hematoxylin and AEC signal); 2. Alignment on hematoxylin images; 3. Creation of transformation matrix on AEC images; 4. For a part of the staining (Edac, Vim, Ki67) an automatic threshold using MaxEntropy was done to remove background, for the rest of the stainings (p16, Krt5, Krt8, Ncad) different threshold was determined using control cell signal (cf. Computational part). Each staining was colored as desired. To further analysis, the composite image was transferred back to QuPath. By hand, the different structures of the gland/tumors were annotated (duct, stroma, juxta-lesion or juxta-tumoral duct, lesion, tumor). To identify all the cells, we used the ‘cell detection’ function based on hematoxylin nucleus staining. We then used the ‘show detection measurement’ function to export the annotation and the intensity signal for all staining for each cell and analyzed it in R.

### Multiplex histological data analysis strategy

The resulting measurements were exported and analyzed in R (4.1.1). Briefly, high signal channels, corresponding to Ki67, Vim were thresholded by the Maximum Entropy algorithm, whereas the remaining channel markers were subjected to a custom thresholding approach. To identify true positive cells for each marker, mean “Cell” signal values were binarized as follows: - non-zero values of the Max Entropy thresholded markers were set to 1, whereas zero values were set to 0. To determine positive cells for p16, Ncad and Krt5, the local minimum after the highest peak was fitted on the density distribution of the merged cells from all the samples corresponding to each marker. Different thresholds were defined for each sample for the following markers: Krt8 and Ecad. Briefly, the “approxfunc” r interpolation function was applied on the density distribution of each marker on each sample, followed by an optimization step using the “optimize” r function to retrieve the local minimum within the interval of the density function. Higher values as compared to each threshold were set to 1, whereas smaller values were set to 0. basic r functions were used to calculate the percentages of positive cells for each or double positive for many markers, and the ggplot package was used for graphical representations. Stromal cells were excluded in the analyses.

### Mammary gland / tumor dissociation and flow cytometry

Samples were cut roughly with dissecting scissors and then with 2 scalpels for approximative 10 min. Then single cell dissociation was done by enzymatic digestion with 3mg/ml collagenase I (Roche, 11088793001) and 100U/ml hyaluronidase (Sigma-Aldrich, H3506) in complete media (HBSS (24020117), 5% SVF) during 1h30 under agitation at 170 rpm at 37°C. Cells were then dissociated in PBS 0,25% Trypsin-Versen (Thermo Fisher Scientific, 15040-033) prewarmed at 37°C for 1min30s with pipetting for 45s. The cell suspension was then treated with dispase 5 mg/ml (Sigma-Aldrich, D4693) and DNase 0,1 mg/ml (Roche, 11284932001) in complete media for 5 min at 37°C. A treatment with Red Blood cell lysis buffer (Thermo Fisher Scientific, 00-4333-57) was carried out then the suspension was filtered at 40µM before counting and FACS staining. Cell suspensions were stained 20 min in dark at 4°C with anti-CD45-APC 1:100 (BioLegend, 103112), anti-CD31-APC 1:100 (BioLegend, 102510), anti-CD24-BV421 1:50 (BioLegend, 101826), anti-CD49f-PE 1:50 (BioLegend, 313622). Cells were resuspended in cytometry media (PBS, BSA, EDTA). For the mammary gland samples, we either recovered the total epithelium or the luminal and basal cells populations separately.

### Single-cell RNA-seq

In accordance with the protocol of 10X Chromium manufacture, the cells were resuspended in PBS 0,04% BSA. Depending on the samples, approximately 3000 or 4000 cells were loaded on the Chromium Single Cell Controller Instrument (Chromium single cell 3’ v3 or 3’ NextGem, 10X Genomics, PN-1000075) in accordance with the manufacturer’s protocol. Libraries were prepared according to the same protocol.

### Bulk and single-cell ChIP-seq

ChIP experiments were performed as previously described (Marsolier et al. 2022) using an anti-H3K27me3 antibody (Cell Signaling Technology, 9733 - C36B11). Bulk sequencing libraries were prepared using the NEBNext Ultra II DNA Library Prep Kit (NEB, E7645S) according to the manufacturer’s instructions. For single-cell experiments, cells were encapsulated on a custom microfluidic device as described before (Grosselin et al. 2019). Cells were stained with DAPI 3µM or with 1µM CFSE during 15 min (CellTrace CFSE, ThermoFisher Scientific, Ref: C34554).

## COMPUTATIONAL ANALYSIS

Code related to the following sections will be deposited on Github (https://github.com/vallotlab).

### Chromium 10X scRNAseq data pre-processing

scRNAseq data acquisition was performed using the 10X toolkit. Briefly, the CellRanger Software Suite (version 3.0.1) was used for demultiplexing, cell barcode assignment and further UMI (Unique molecular Identifier) quantification. The pre-built mm10 reference genome proposed by 10X Genomics ((https://support.10xgenomics.com/single-cell-gene-expression/software/downloads/latest) was used to align the reads. All the *in vivo* mouse datasets were analyzed together, without performing any batch correction. Doublet removal step was included in the 10X workflow, and was performed by the “emptyDrops” function from DropletUtils at an FDR of 0.01.

### Quality Control (QC) for scRNAseq data analysis

Low quality cells were defined as having aberrant values for the type and number of genes/UMIs detected. We evaluated the distribution of the total number of genes, molecules (UMIs) and the fraction of UMIs mapped to mitochondrial (MT) genes and set up thresholds to filter out those cells. Three upper cutoffs of 30% UMIs mapped to MT genes, 10,000 genes and 100,000 nUMIs were fixed to get rid of outliers. Additionally, cells with less than 1000 detected genes were excluded. This resulted in a total of 17,330 high quality cells, which were used for further analysis.

### scRNAseq data Normalization

Normalization and variance stabilization were performed using the SCTransform method, implemented in the “SCTransform’’ function from the Seurat Suit. SCTransform omits the need for heuristic steps comprising log-transformation and pseudo-count addition, and results in improved downstream analytical steps. More recently, SCTransform also supports using the <u>glmGamPoi</u> package. Briefly, this method fits a “Gamma-Poisson Generalized Linear Model” to the overdispersed count matrices due to the high sparsity of the scRNAseq data, and results in a substantial improvement of the variance stabilization.

### scRNAseq data dimension reduction and clustering

Principal Component Analysis (PCA) was performed on the top 3000 Highly variable genes of the SCT assay from the SCTransform step, to reduce the data dimensionality. The top 60 PCs were further used to perform graph-based clustering and community (cell cluster) detection.

All the Uniform Manifold Approximation and Projection (UMAP) plots were computed using the “RunUMAP” Seurat function with default parameters (“uwot” as umap.method, n.neighbours=30, distance metric= “cosine”, min.dist=0.3) and “random.state=42”. The two-dimensional UMAP coordinates were calculated using the top 60 PCs previously computed on the SCT assay. For the sake of clarity, once the epithelial compartment is sub clustered, the same UMAP embeddings were used to represent the “transitioning cell clusters”. Further “zoom ins” were performed using the corresponding umap coordinates of the cells of interest.

### Graph-based clustering and cell cluster identification

Cell clustering was performed using a two-step wise approach, using the “FIndNeighbours’’ and “FindClusters’’ respectively. Briefly, a k-Nearest Neighbours (kNN) graph is built on the dissimilarity matrix based on the pairwise euclidean distance between cells in the PCA space (using the previously computed 60 PCs). Edges are drawn between nodes (cells) with similar expression patterns (Jaccard Similarity). Edge weights are refined based on their shared overlap in their neighborhood.

“FindClusters” function was used to cluster the cells, using the Louvain algorithm as default, setting the resolution parameter to 1.2 to ensure an optimal granularity and stability of the cell clusters.

### scRNAseq cluster annotation

Manual annotation of the cell clusters was performed on the merged samples on a two-steps basis. First, the cell clusters were annotated according to the major cell compartments, using well-established canonical markers. The latter included: Immune (*Ptprc+, Cd68+, Cd52+*), Epithelial (*Epcam, Krt5, Krt8, Elf5*), Endothelial (*Pecam1, Fabp4, Apold1*) and Fibroblasts (*Mgp, Dpep1, Col3a1*). Briefly, we computed the mean expression of each gene across the cells belonging to each cluster, to obtain a pseudo-bulked matrix containing only the genes of interest. A classical hierarchical clustering was performed on the clusters based on their correlation distance matrix to determine the cell cluster groups “Metaclusters” which displayed similar expression levels for each canonical gene signature. According to the dendrogram length, computed using the “ward.D” method, 5 meta-clusters were identified. Each meta-cluster was assigned the cell type name for which the canonical genes were mostly expressed, as compared to the remaining genes. For instance, COl3a1 displayed the highest expression level in the meta-cluster “1”. Therefore, all cell clusters previously defined (see **Graph-based clustering and cell cluster identification** section above) belonging to metacluster “1” are labelled as “Fibroblasts”.

### Refined Epithelial clusters annotation

The epithelial compartment was further sub clustered to better explore the cell subtype/state heterogeneity between the control and the tumor samples. SCTrasnform, PCA dimension reduction and clustering steps were run on the subsetted clusters of interest. To achieve a high-resolutive cell subtype annotation, DIfferential expression (DE) was performed using the “FindAllMarkers” function. Briefly, a non-parametric Wilcoxon-Sum rank test was performed on a “1 cluster vs all” basis, setting a log2 Fold Change (FC) threshold at 0.5, and keeping only genes expressed in at least 30% of the cell clusters (to ensure expression homogeneity within the cluster). Associated p-values were corrected using the “Bonferroni” correction method, with a set threshold at 5%.

An automated function was designed to annotate the clusters. It takes as input the top 10 logFC ranked geneset for each cluster, and initially computes the contribution percentage of each tumor size feature of our dataset (control, control with primary lesions, small tumor (ST), medium tumor (MT) and large tumor (LT)) to each cell cluster. For a given cluster, if the major contributor is the control dataset, the function intersects the corresponding top genes with a knowledge-driven gene list of the known epithelial cell types (including basal, luminal progenitor, alveolar-differentiated, hormone-sensing …) and labels the cluster with the corresponding cell type. If most of the cells (> 60%) were from tumor samples, the subtype name would be the concatenation of the top gene name with the tumor size symbol (ST, MT or LT).

### Differential expression

Differential gene expression (DGE) analysis was performed using “FindMarkers” function. Non parametric Wilcoxon sum rank test was used to identify genes with an abs(FC)> 0.5 at an FDR of 0.05. To ensure cell cluster homogeneity, we set a lower cutoff of 30% of cells expressing a given gene.

### Pathway Enrichment Analysis (PEA)

Pathway Enrichment Analysis was performed on the significantly differentially expressed gene lists using the Hallmark collection from the Molecular Signature Database (MSigDB). The latter was loaded into the R session using the “msigdbr” package available on Bioconductor. Gene Set Enrichment Analysis was performed using the “enricher” function from the “msigdbr” package. Only significantly enriched pathways (adjusted P-values < 0.05) were considered.

### Signature construction

Transcriptional signatures were constructed from the gene lists contributing to each corresponding enriched pathway, using the “AUCell” package available on Github ( ). Briefly, the genes of a given cell vs.gene data matrix are ranked based on their expression levels in each cell. UCell computes then a Mann-Whitney U statistic (which is similar to AUC Area Under Curve), which is further used to evaluate gene signatures on the gene expression ranks of individual cells. We computed the gene signatures using the wrapper function “AddModuleScore_UCell”, giving as input a list of features, along with the seurat object.

### Trajectory inference - Slingshot

Pseudotime ordering of cells was conducted using Slingshot (Github link), with default parameters, giving as input the UMAP coordinates and setting the starting cluster as the luminal progenitors “LP”, with stretch=2.

To ease the interpretation of the trajectory, we performed SLingshot only on the transitioning compartment, including (“LP”, Alveolar differentiated “Avd”, Luminal differentiated hormone-sensing “Luminal H-S”, and the annotated clusters of the small tumor. Downstream analytical steps were performed only on the longest branch starting from the “LP” and ending in the “Fgf8+ ST” cluster.

### Contribution of genes to a branch tree

The aim of this section was to identify the most contributing genes to the transition observed from the Slingshot trajectory inference. To do so, a cell vs.gene expression matrix was created including the contributing cells to the longest branch, and the top 2000 highly variable genes. We then applied a random forest regression model using 500 trees to predict the genes which contribute the most to predict pseudotime values (the response variable). The features (genes) were sorted according to their “variable.importance”parameter after the model was fit.

### Associated pathways to pseudotime values prediction

We computed the mean expression values of the selected top 200 most important predictive genes to get pseudo-bulked matrices for the transitioning cells. To cluster the genes according to their profile correlation with pseudotime values, a pairwise-correlation matrix, followed by a hierarchical clustering were performed. 5 gene groups were obtained, each having a distinct profile along pseudotime. PEA (see below) was performed on each gene set, followed by a signature construction step and ultimately visualized on the UMAP embeddings.

### Partition-based graph abstraction (PAGA)

PAGA was performed using “scanpy” Python library loaded on RStudio using “reticulate” R package. Default parameters were used to construct the graph partition, and a threshold of 0.1 was set to preserve the highly connected nodes. Connectivity scores were extracted from the PAGA output, along with the nodes and edges connections. Centrality scores (number of edges) were computed by counting the number of edges that passed the cutoff (0.15) for each cell cluster.

### Potential of Heat-diffusion for Affinity-based Transition Embedding (PHATE)

PHATE was used as a visualization method to investigate continual progressions, branches and clusters in our data. Briefly, PHATE uses an information-geometric distance between cells (data points) to capture both local and global nonlinear structures, setting knn = 20, t (diffusion parameter) =40 as input parameters.

### Copy Number Variation (CNV) inference from scRNAseq data

CNVs were inferred using inferCNV (https://github.com/broadinstitute/infercnv) with default parameters, taking as reference the basal cells. We extracted residual cell matrices, binarized the values using the 10th as lower and 90th percentile as higher thresholds, to get -1 (if the value < 10th percentile); +1 (if the value is higher than the 90th percentile) and 0 if the value is in between the two thresholds. To estimate the percentage of altered genome, we calculated the absolute value of binarized matrices, and counted the number of 0s and 1s aggregated by chromosome. These values were added to the metadata of the scRNAseq Seurat object.

### TCGA_Breast cancer dataset

To compare the expression levels of CDKN2A, P16-signature, EMT and apoptosis pathway signatures, between non-diseased healthy tissues, tumor-adjacent normal tissue and tumor breast tissues, we harnessed breast tissues datasets from TCGA and GTEx consortia from normalized transcriptomic data available from Github (https://github.com/mskcc/RNAseqDB/tree/master/data/normalized). We constructed the gene signatures using the UCell package, and compared the tissue types using Wilcoxon T tests.

### scRNAseq data analysis of normal, preneoplastic and tumorigenic states in the human breast

We downloaded the dataset from GEO, using the accession number: GSE161529. Briefly, we selected only the normal epithelium samples from pre-menopausal women (n=6), tumor samples (labeled as Triple Negative tumor, and Triple negative (Brca) tumor) (ntotal=8), and the nulliparous, pre-menopausal pre-neoplastic Brca1 samples (n=2). After sample merging, SCT normalization, dimension reduction and graph-based clustering, we selected the cell clusters expressing epithelial markers (Epcam, Krt8, Krt5) for further analysis. The same procedure was conducted on the epithelial compartment, followed by a finer annotation of the cell clusters using canonical markers of epithelial sub-populations. To point out the epithelial population which underwent major transcriptional modifications upon Brca1 deficiency as compared to the normal population, we subset the luminal progenitor (LP), Basal and mature luminal (ML) clusters. For each subpopulation, principal component analysis (PCA) was performed, and the top 20 variable PCs were kept. To identify the main PC drivers of a normal/preneoplastic gradient, we tested whether the cell distributions along each PC coordinate were the same, using a Kosmogorov Smirnov nonparametric test. We selected the PCs with a significant p-value (<0.05) and a D-value > quantile(D-value,0.8). Alternatively, a linear regression method was tested to select the top predictive PCs to separate cells labeled as preneoplastic from normal ones. Both methods indicated similar PCs. Next, to identify the epithelial sub-population for which the PCs were the most discriminant, we ranked the top “informative” PCs according to their percentage of variance explained. Pathway enrichment analysis was performed on the top genes (ranked by eigenvalues) which contributed most to the PC part corresponding to preneoplastic cells.

## DATA AVAILABILITY

The datasets described in this study have been deposited in the prive GEO repository GSE200444, accessible with the token gtotisgiftopnqr.

## ACKNOWLEDGMENTS

We thank Dr S. Fre for providing critical discussion. We also thank the animal facility, the sequencing and imaging platforms from Institut Curie. We thank Dr J. Jonkers for providing mouse strains.

## REFERENCES

Aird, Katherine M., and Rugang Zhang. 2013. “Detection of Senescence-Associated Heterochromatin Foci (SAHF).” Methods in Molecular Biology 965: 185–96.

Ansieau, Stéphane, Jeremy Bastid, Agnès Doreau, Anne-Pierre Morel, Benjamin P. Bouchet, Clémence Thomas, Frédérique Fauvet, et al. 2008. “Induction of EMT by Twist Proteins as a Collateral Effect of Tumor-Promoting Inactivation of Premature Senescence.” Cancer Cell 14 (1): 79–89.

Bach, Karsten, Sara Pensa, Marta Grzelak, James Hadfield, David J. Adams, John C. Marioni, and Walid T. Khaled. 2017. “Differentiation Dynamics of Mammary Epithelial Cells Revealed by Single-Cell RNA Sequencing.” Nature Communications 8 (1): 2128.

Bach, Karsten, Sara Pensa, Marija Zarocsinceva, Katarzyna Kania, Julie Stockis, Silvain Pinaud, Kyren A. Lazarus, et al. 2021. “Time-Resolved Single-Cell Analysis of Brca1 Associated Mammary Tumourigenesis Reveals Aberrant Differentiation of Luminal Progenitors.” Nature Communications 12 (1): 1502.

Bankhead, Peter, Maurice B. Loughrey, José A. Fernández, Yvonne Dombrowski, Darragh G. McArt, Philip D. Dunne, Stephen McQuaid, et al. n.d. “QuPath: Open Source Software for Digital Pathology Image Analysis.” https://doi.org/10.1101/099796.

Berger, Ashton C., Anil Korkut, Rupa S. Kanchi, Apurva M. Hegde, Walter Lenoir, Wenbin Liu, Yuexin Liu, et al. 2018. “A Comprehensive Pan-Cancer Molecular Study of Gynecologic and Breast Cancers.” Cancer Cell 33 (4): 690–705.e9.

Bianchini, Giampaolo, Justin M. Balko, Ingrid A. Mayer, Melinda E. Sanders, and Luca Gianni. 2016. “Triple-Negative Breast Cancer: Challenges and Opportunities of a Heterogeneous Disease.” Nature Reviews. Clinical Oncology 13 (11): 674–90.

Bizhanova, Aizhan, and Paul D. Kaufman. 2021. “Close to the Edge: Heterochromatin at the Nucleolar and Nuclear Peripheries.” Biochimica et Biophysica Acta (BBA) - Gene Regulatory Mechanisms. https://doi.org/10.1016/j.bbagrm.2020.194666.

Buj, Raquel, Kelly E. Leon, Marlyn A. Anguelov, and Katherine M. Aird. 2021. “Suppression of p16 Alleviates the Senescence-Associated Secretory Phenotype.” Aging 13 (3): 3290–3312.

Campisi, Judith, and Fabrizio d’Adda di Fagagna. 2007. “Cellular Senescence: When Bad Things Happen to Good Cells.” Nature Reviews. Molecular Cell Biology 8 (9): 729–40.

Cancer Genome Atlas Network. 2012. “Comprehensive Molecular Portraits of Human Breast Tumours.” Nature 490 (7418): 61–70.

Cao, Liu, Wenmei Li, Sangsoo Kim, Steven G. Brodie, and Chu-Xia Deng. 2003. “Senescence, Aging, and Malignant Transformation Mediated by p53 in Mice Lacking the Brca1 Full-Length Isoform.” Genes & Development 17 (2): 201–13.

Chakrabarti, Rumela, Julie Hwang, Mario Andres Blanco, Yong Wei, Martin Lukačišin, Rose-Anne Romano, Kirsten Smalley, et al. 2012. “Elf5 Inhibits the Epithelial–mesenchymal Transition in Mammary Gland Development and Breast Cancer Metastasis by Transcriptionally Repressing Snail2.” Nature Cell Biology. https://doi.org/10.1038/ncb2607.

Chaligné, Ronan, Tatiana Popova, Marco-Antonio Mendoza-Parra, Mohamed-Ashick M. Saleem, David Gentien, Kristen Ban, Tristan Piolot, et al. 2015. “The Inactive X Chromosome Is Epigenetically Unstable and Transcriptionally Labile in Breast Cancer.” Genome Research 25 (4): 488–503.

Cmarko, D., J. Smigova, L. Minichova, and A. Popov. 2008. “Nucleolus: The Ribosome Factory.” Histology and Histopathology 23 (10): 1291–98.

Collado, Manuel, and Manuel Serrano. 2010. “Senescence in Tumours: Evidence from Mice and Humans.” Nature Reviews Cancer. https://doi.org/10.1038/nrc2772.

Cristea, Simona, and Kornelia Polyak. 2018. “Dissecting the Mammary Gland One Cell at a Time.” Nature Communications. https://doi.org/10.1038/s41467-018-04905-2.

Di Micco, Raffaella, Valery Krizhanovsky, Darren Baker, and Fabrizio d’Adda di Fagagna. 2021. “Cellular Senescence in Ageing: From Mechanisms to Therapeutic Opportunities.” Nature Reviews. Molecular Cell Biology 22 (2): 75–95.

Engebraaten, Olav, Hans Kristian Moen Vollan, and Anne-Lise Børresen-Dale. 2013. “Triple-Negative Breast Cancer and the Need for New Therapeutic Targets.” The American Journal of Pathology. https://doi.org/10.1016/j.ajpath.2013.05.033.

Ewald, Jonathan A., Joshua A. Desotelle, George Wilding, and David F. Jarrard. 2010. “Therapy-Induced Senescence in Cancer.” Journal of the National Cancer Institute 102 (20): 1536–46.

Fitsiou, Eleni, Abel Soto-Gamez, and Marco Demaria. 2021. “Biological Functions of Therapy-Induced Senescence in Cancer.” Seminars in Cancer Biology. https://doi.org/10.1016/j.semcancer.2021.03.021.

Fridman, A. L., and M. A. Tainsky. 2008. “Critical Pathways in Cellular Senescence and Immortalization Revealed by Gene Expression Profiling.” Oncogene 27 (46): 5975–87.

Ganesan, Shridar, Daniel P. Silver, Ronny Drapkin, Roger Greenberg, Jean Feunteun, and David M. Livingston. 2004. “Association of BRCA1 with the Inactive X Chromosome and XIST RNA.” Philosophical Transactions of the Royal Society of London. Series B: Biological Sciences. https://doi.org/10.1098/rstb.2003.1371.

Gao, Ruli, Alexander Davis, Thomas O. McDonald, Emi Sei, Xiuqing Shi, Yong Wang, Pei-Ching Tsai, et al. 2016. “Punctuated Copy Number Evolution and Clonal Stasis in Triple-Negative Breast Cancer.” Nature Genetics 48 (10): 1119–30.

Gray, G. Kenneth, Carman Man-Chung Li, Jennifer M. Rosenbluth, Laura M. Selfors, Nomeda Girnius, Jia-Ren Lin, Ron C. J. Schackmann, et al. 2022. “A Human Breast Atlas Integrating Single-Cell Proteomics and Transcriptomics.” *Developmental Cell*, May. https://doi.org/10.1016/j.devcel.2022.05.003.

Grosselin, Kevin, Adeline Durand, Justine Marsolier, Adeline Poitou, Elisabetta Marangoni, Fariba Nemati, Ahmed Dahmani, et al. 2019. “High-Throughput Single-Cell ChIP-Seq Identifies Heterogeneity of Chromatin States in Breast Cancer.” Nature Genetics 51 (6): 1060–66.

Hanahan, Douglas, and Robert A. Weinberg. 2016. “The Hallmarks of Cancer.” Oxford Textbook of Oncology. https://doi.org/10.1093/med/9780199656103.003.0001.

Harada, Takamasa, Joe Swift, Jerome Irianto, Jae-Won Shin, Kyle R. Spinler, Avathamsa Athirasala, Rocky Diegmiller, P. C. Dave P. Dingal, Irena L. Ivanovska, and Dennis E. Discher. 2014. “Nuclear Lamin Stiffness Is a Barrier to 3D Migration, but Softness Can Limit Survival.” The Journal of Cell Biology 204 (5): 669–82.

Herranz, Nicolás, and Jesús Gil. 2018. “Mechanisms and Functions of Cellular Senescence.” Journal of Clinical Investigation. https://doi.org/10.1172/jci95148.

Ito, Takahiro, Yee Voan Teo, Shane A. Evans, Nicola Neretti, and John M. Sedivy. 2018. “Regulation of Cellular Senescence by Polycomb Chromatin Modifiers through Distinct DNA Damage- and Histone Methylation-Dependent Pathways.” Cell Reports 22 (13): 3480–92.

Jonkers, Jos, Ralph Meuwissen, Hanneke van der Gulden, Hans Peterse, Martin van der Valk, and Anton Berns. 2001. “Synergistic Tumor Suppressor Activity of BRCA2 and p53 in a Conditional Mouse Model for Breast Cancer.” Nature Genetics. https://doi.org/10.1038/ng747.

Koppelstaetter, Christian, Gabriele Schratzberger, Paul Perco, Johannes Hofer, Walter Mark, Robert Ollinger, Rainer Oberbauer, et al. 2008. “Markers of Cellular Senescence in Zero Hour Biopsies Predict Outcome in Renal Transplantation.” Aging Cell 7 (4): 491–97.

Kristiansen, M., G. P. S. Knudsen, P. Maguire, S. Margolin, J. Pedersen, A. Lindblom, and K. H. Ørstavik. 2005. “High Incidence of Skewed X Chromosome Inactivation in Young Patients with Familial Non-BRCA1/BRCA2 Breast Cancer.” Journal of Medical Genetics 42 (11): 877–80.

Lamouille, Samy, Jian Xu, and Rik Derynck. 2014. “Molecular Mechanisms of Epithelial-Mesenchymal Transition.” Nature Reviews. Molecular Cell Biology 15 (3): 178–96.

Liberzon, Arthur, Chet Birger, Helga Thorvaldsdóttir, Mahmoud Ghandi, Jill P. Mesirov, and Pablo Tamayo. 2015. “The Molecular Signatures Database Hallmark Gene Set Collection.” Cell Systems. https://doi.org/10.1016/j.cels.2015.12.004.

Lim, Elgene, François Vaillant, Di Wu, Natasha C. Forrest, Bhupinder Pal, Adam H. Hart, Marie-Liesse Asselin-Labat, et al. 2009. “Aberrant Luminal Progenitors as the Candidate Target Population for Basal Tumor Development in BRCA1 Mutation Carriers.” Nature Medicine 15 (8): 907–13.

Lim, Su Bin, Swee Jin Tan, L. I. M. Wan-Teck, and Chwee Teck Lim. 2017. “An Extracellular Matrix-Related Prognostic and Predictive Indicator for Early-Stage Non-Small Cell Lung Cancer.” Nature Communications. https://doi.org/10.1038/s41467-017-01430-6.

Liu, Xiaoling, Henne Holstege, Hanneke van der Gulden, Marcelle Treur-Mulder, John Zevenhoven, Arno Velds, Ron M. Kerkhoven, et al. 2007. “Somatic Loss of BRCA1 and p53 in Mice Induces Mammary Tumors with Features of Human BRCA1-Mutated Basal-like Breast Cancer.” Proceedings of the National Academy of Sciences of the United States of America 104 (29): 12111–16.

Marra, Antonio, Dario Trapani, Giulia Viale, Carmen Criscitiello, and Giuseppe Curigliano. 2020. “Practical Classification of Triple-Negative Breast Cancer: Intratumoral Heterogeneity, Mechanisms of Drug Resistance, and Novel Therapies.” NPJ Breast Cancer 6 (October): 54.

Marsolier, Justine, Pacôme Prompsy, Adeline Durand, Anne-Marie Lyne, Camille Landragin, Amandine Trouchet, Sabrina Tenreira Bento, et al. 2022. “H3K27me3 Conditions Chemotolerance in Triple-Negative Breast Cancer.” Nature Genetics 54 (4): 459–68.

Martins, Filipe C., Subhajyoti De, Vanessa Almendro, Mithat Gönen, So Yeon Park, Joanne L. Blum, William Herlihy, et al. 2012. “Evolutionary Pathways in BRCA1-Associated Breast Tumors.” Cancer Discovery 2 (6): 503–11.

Milanovic, Maja, Dorothy N. Y. Fan, Dimitri Belenki, J. Henry M. Däbritz, Zhen Zhao, Yong Yu, Jan R. Dörr, et al. 2018. “Senescence-Associated Reprogramming Promotes Cancer Stemness.” Nature 553 (7686): 96–100.

Molyneux, Gemma, Felipe C. Geyer, Fiona-Ann Magnay, Afshan McCarthy, Howard Kendrick, Rachael Natrajan, Alan MacKay, et al. 2010. “BRCA1 Basal-like Breast Cancers Originate from Luminal Epithelial Progenitors and Not from Basal Stem Cells.” Cell Stem Cell. https://doi.org/10.1016/j.stem.2010.07.010.

Moon, Kevin R., David van Dijk, Zheng Wang, Scott Gigante, Daniel B. Burkhardt, William S. Chen, Kristina Yim, et al. 2019. “Visualizing Structure and Transitions in High-Dimensional Biological Data.” Nature Biotechnology 37 (12): 1482–92.

Nallanthighal, Sameera, James Patrick Heiserman, and Dong-Joo Cheon. 2021. “Collagen Type XI Alpha 1 (COL11A1): A Novel Biomarker and a Key Player in Cancer.” Cancers. https://doi.org/10.3390/cancers13050935.

Nieto, M. Angela, M. Angela Nieto, Ruby Yun-Ju Huang, Rebecca A. Jackson, and Jean Paul Thiery. 2016. “EMT: 2016.” Cell. https://doi.org/10.1016/j.cell.2016.06.028.

Onitilo, Adedayo A., Jessica M. Engel, Robert T. Greenlee, and Bickol N. Mukesh. 2009. “Breast Cancer Subtypes Based on ER/PR and Her2 Expression: Comparison of Clinicopathologic Features and Survival.” Clinical Medicine & Research 7 (1-2): 4–13.

Pal, Bhupinder, Toula Bouras, Wei Shi, François Vaillant, Julie M. Sheridan, Naiyang Fu, Kelsey Breslin, et al. 2013. “Global Changes in the Mammary Epigenome Are Induced by Hormonal Cues and Coordinated by Ezh2.” Cell Reports 3 (2): 411–26.

Pal, Bhupinder, Yunshun Chen, François Vaillant, Bianca D. Capaldo, Rachel Joyce, Xiaoyu Song, Vanessa L. Bryant, et al. 2021. “A Single-Cell RNA Expression Atlas of Normal, Preneoplastic and Tumorigenic States in the Human Breast.” The EMBO Journal 40 (11): e107333.

Paluvai, Harikrishnareddy, Eros Di Giorgio, and Claudio Brancolini. 2020. “The Histone Code of Senescence.” Cells. https://doi.org/10.3390/cells9020466.

Pastushenko, Ievgenia, Audrey Brisebarre, Alejandro Sifrim, Marco Fioramonti, Tatiana Revenco, Soufiane Boumahdi, Alexandra Van Keymeulen, et al. 2018. “Identification of the Tumour Transition States Occurring during EMT.” Nature 556 (7702): 463–68.

Patel, A. P., I. Tirosh, J. J. Trombetta, A. K. Shalek, S. M. Gillespie, H. Wakimoto, D. P. Cahill, et al. 2014. “Single-Cell RNA-Seq Highlights Intratumoral Heterogeneity in Primary Glioblastoma.” Science. https://doi.org/10.1126/science.1254257.

Pearce, Oliver M. T., Robin M. Delaine-Smith, Eleni Maniati, Sam Nichols, Jun Wang, Steffen Böhm, Vinothini Rajeeve, et al. 2018. “Deconstruction of a Metastatic Tumor Microenvironment Reveals a Common Matrix Response in Human Cancers.” Cancer Discovery. https://doi.org/10.1158/2159-8290.cd-17-0284.

Pervolarakis, Nicholas, Quy H. Nguyen, Justice Williams, Yanwen Gong, Guadalupe Gutierrez, Peng Sun, Darisha Jhutty, et al. 2020. “Integrated Single-Cell Transcriptomics and Chromatin Accessibility Analysis Reveals Regulators of Mammary Epithelial Cell Identity.” Cell Reports 33 (3): 108273.

Polak, Paz, Jaegil Kim, Lior Z. Braunstein, Rosa Karlic, Nicholas J. Haradhavala, Grace Tiao, Daniel Rosebrock, et al. 2017. “A Mutational Signature Reveals Alterations Underlying Deficient Homologous Recombination Repair in Breast Cancer.” Nature Genetics 49 (10): 1476–86.

Probst, Aline V., and Geneviève Almouzni. 2008. “Pericentric Heterochromatin: Dynamic Organization during Early Development in Mammals.” Differentiation. https://doi.org/10.1111/j.1432-0436.2007.00220.x.

Remark, Romain, Taha Merghoub, Niels Grabe, Geert Litjens, Diane Damotte, Jedd D. Wolchok, Miriam Merad, and Sacha Gnjatic. 2016. “In-Depth Tissue Profiling Using Multiplexed Immunohistochemical Consecutive Staining on Single Slide.” Science Immunology. https://doi.org/10.1126/sciimmunol.aaf6925.

Roupakia, Eugenia, Georgios S. Markopoulos, and Evangelos Kolettas. 2021. “Genes and Pathways Involved in Senescence Bypass Identified by Functional Genetic Screens.” Mechanisms of Ageing and Development 194 (March): 111432.

Scully, R., and D. M. Livingston. 2000. “In Search of the Tumour-Suppressor Functions of BRCA1 and BRCA2.” Nature 408 (6811): 429–32.

Sedic, Maja, Adam Skibinski, Nelson Brown, Mercedes Gallardo, Peter Mulligan, Paula Martinez, Patricia J. Keller, et al. 2015. “Haploinsufficiency for BRCA1 Leads to Cell-Type-Specific Genomic Instability and Premature Senescence.” Nature Communications 6 (June): 7505.

Selbert, S., D. J. Bentley, D. W. Melton, D. Rannie, P. Lourenço, C. J. Watson, and A. R. Clarke. 1998. “Efficient BLG-Cre Mediated Gene Deletion in the Mammary Gland.” Transgenic Research 7 (5): 387–96.

Severino, V., N. Alessio, A. Farina, A. Sandomenico, M. Cipollaro, G. Peluso, U. Galderisi, and A. Chambery. 2013. “Insulin-like Growth Factor Binding Proteins 4 and 7 Released by Senescent Cells Promote Premature Senescence in Mesenchymal Stem Cells.” Cell Death & Disease 4 (November): e911.

Sirchia, Silvia M., Lisetta Ramoscelli, Francesca R. Grati, Floriana Barbera, Danila Coradini, Franca Rossella, Giovanni Porta, et al. 2005. “Loss of the Inactive X Chromosome and Replication of the Active X in BRCA1-Defective and Wild-Type Breast Cancer Cells.” Cancer Research 65 (6): 2139–46.

Stefansson, Olafur Andri, Jon Gunnlaugur Jonasson, Kristrun Olafsdottir, Holmfridur Hilmarsdottir, Gudridur Olafsdottir, Manel Esteller, Oskar Thor Johannsson, and Jorunn Erla Eyfjord. 2011. “CpG Island Hypermethylation of BRCA1 and Loss of pRb as Co-Occurring Events in Basal/triple-Negative Breast Cancer.” Epigenetics: Official Journal of the DNA Methylation Society 6 (5): 638–49.

Stone, Caroline, Nuala McCabe, and Alan Ashworth. 2003. “X-Chromosome Inactivation: X Marks the Spot for BRCA1.” Current Biology: CB 13 (2): R63–64.

Street, Kelly, Davide Risso, Russell B. Fletcher, Diya Das, John Ngai, Nir Yosef, Elizabeth Purdom, and Sandrine Dudoit. 2018. “Slingshot: Cell Lineage and Pseudotime Inference for Single-Cell Transcriptomics.” BMC Genomics 19 (1): 477.

Timms, Kirsten M., Victor Abkevich, Elisha Hughes, Chris Neff, Julia Reid, Brian Morris, Saritha Kalva, et al. 2014. “Association of BRCA1/2defects with Genomic Scores Predictive of DNA Damage Repair Deficiency among Breast Cancer Subtypes.” Breast Cancer Research. https://doi.org/10.1186/s13058-014-0475-x.

Vincent-Salomon, Anne, Carine Ganem-Elbaz, Elodie Manié, Virginie Raynal, Xavier Sastre-Garau, Dominique Stoppa-Lyonnet, Marc-Henri Stern, and Edith Heard. 2007. “X Inactive–Specific Transcript RNA Coating and Genetic Instability of the X Chromosome in *BRCA1* Breast Tumors.” Cancer Research. https://doi.org/10.1158/0008-5472.can-07-0465.

Visvader, Jane E., and John Stingl. 2014. “Mammary Stem Cells and the Differentiation Hierarchy: Current Status and Perspectives.” Genes & Development 28 (11): 1143–58.

Watson, Christine J., and Walid T. Khaled. 2008. “Mammary Development in the Embryo and Adult: A Journey of Morphogenesis and Commitment.” Development 135 (6): 995–1003.

Wolf, F. Alexander, Fiona K. Hamey, Mireya Plass, Jordi Solana, Joakim S. Dahlin, Berthold Göttgens, Nikolaus Rajewsky, Lukas Simon, and Fabian J. Theis. 2019. “PAGA: Graph Abstraction Reconciles Clustering with Trajectory Inference through a Topology Preserving Map of Single Cells.” Genome Biology 20 (1): 59.

Yang, Jing, Parker Antin, Geert Berx, Cédric Blanpain, Thomas Brabletz, Marianne Bronner, Kyra Campbell, et al. 2020. “Guidelines and Definitions for Research on Epithelial-Mesenchymal Transition.” Nature Reviews. Molecular Cell Biology 21 (6): 341–52.

Yochum, Zachary A., Jessica Cades, Lucia Mazzacurati, Neil M. Neumann, Susheel K. Khetarpal, Suman Chatterjee, Hailun Wang, et al. 2017. “A First-in-Class TWIST1 Inhibitor with Activity in Oncogene-Driven Lung Cancer.” Molecular Cancer Research. https://doi.org/10.1158/1541-7786.mcr-17-0298.

Zhang, Yun, Joana Liu Donaher, Sunny Das, Xin Li, Ferenc Reinhardt, Jordan A. Krall, Arthur W. Lambert, et al. 2022. “Genome-Wide CRISPR Screen Identifies PRC2 and KMT2D-COMPASS as Regulators of Distinct EMT Trajectories That Contribute Differentially to Metastasis.” Nature Cell Biology 24 (4): 554–64.

