## SupplementaryFigures for "Luminal progenitors undergo partial epithelial-to-mesenchymal transition at the onset of basal-like breast tumorigenesis"

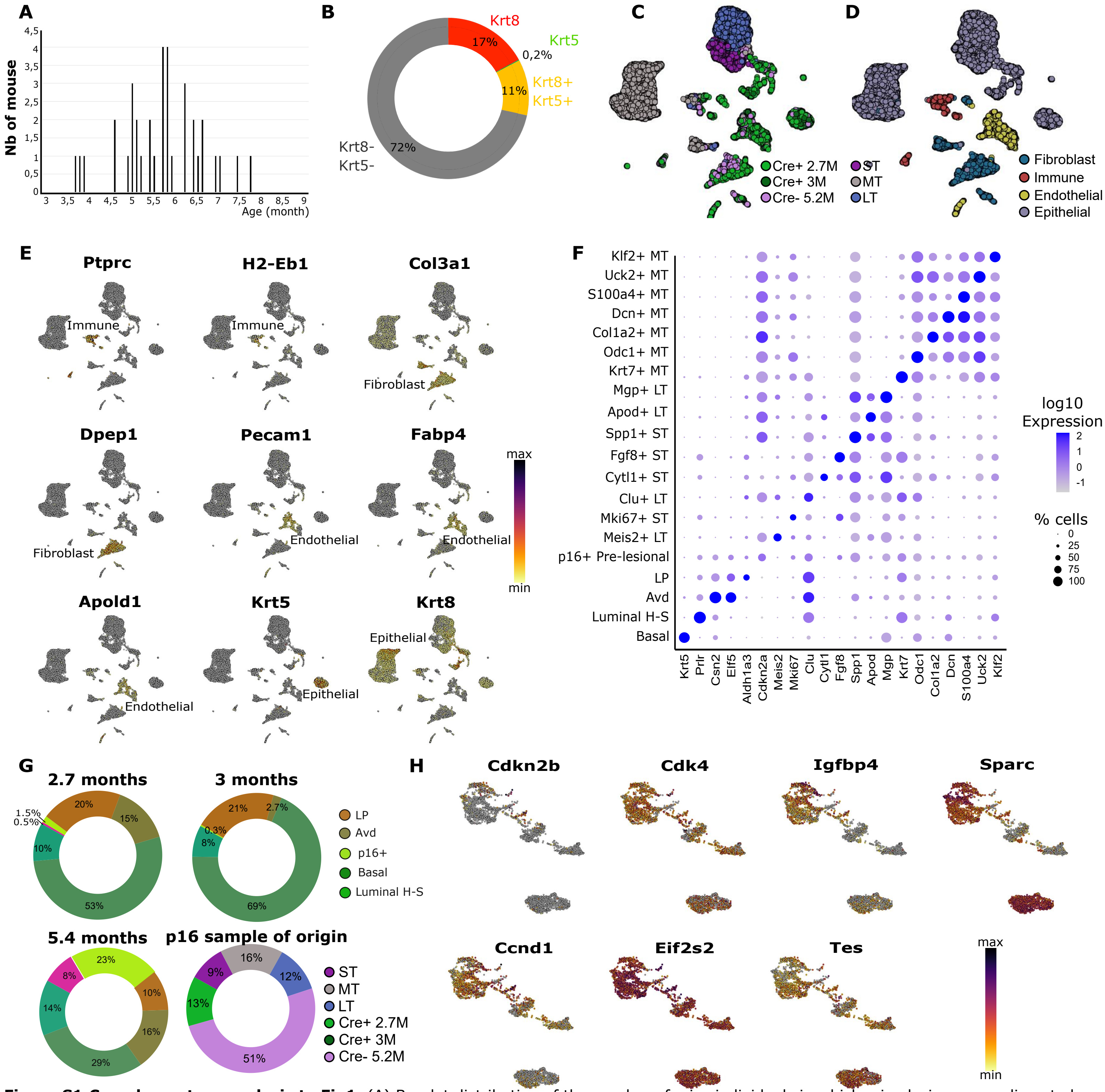


**Figure S1:Complementary analysis to Fig1.** (A) Barplot distribution of the number of mice individuals in which microlesions were dissected; (B) Donut plot representation of the repartition of cells according to Krt8, Krt5 fluorescent stainings;UMAP dimension reduction of the mammary gland compartment landscape; in which cells were colored according to (C) the sample of origin and (D) the corresponding annotated celltype; (E) Umap plots of the mammary gland compartment, colored by expression levels of canonical markers of immune, fibroblast, epithelial and endothelial cell types; (F) Dot plot representation of average expression levels of top expressed genes by each subcluster of epithelial cells. Average expression levels are color-coded, and percentage of cells expressing the genes is size-coded; (G) Donut plot representation of the repartition of epithelial sup-populations between the 3 control samples, profiles using scRNAseq data technology along with the repartition of the samples of origin of the p16 pre-tumoral population; (H) Umap plots of the subset of the epithelial compartment, colored by the expression levels of genes involved in cellular senescence.

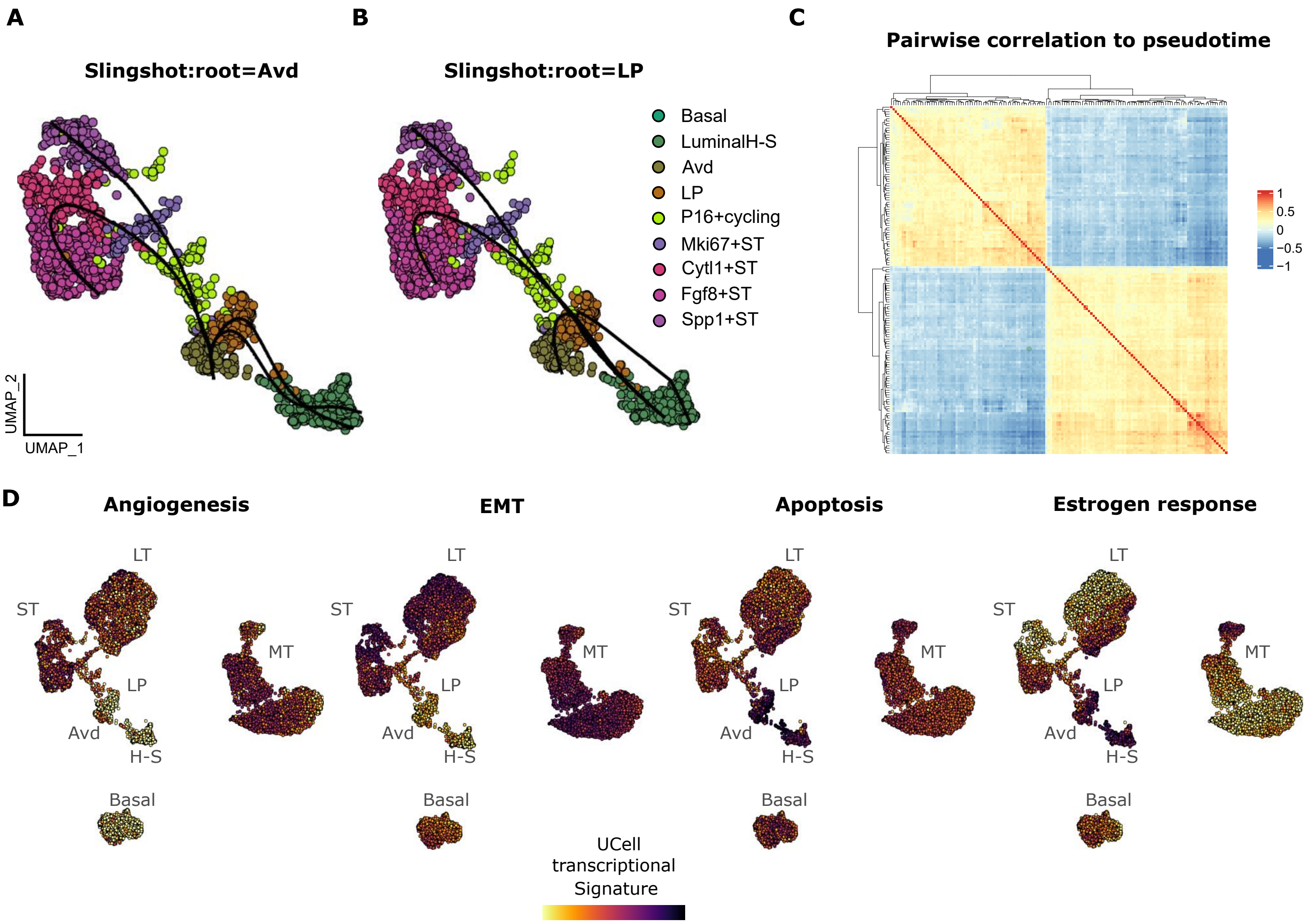


**Figure S2: Characterization of p16 pre-tumoral cell transitioning genes**. Umap plot of the transitioning cell, colored by subtype annotation, on which transition path is shown by the passing-by line on the cells; taking as reference either the LP (A), or Avd (B);
(C) Heatmap representation of pairwise correlation values between pseudotime and the mostly correlated genes (in absolute values) to pseudotome values along the path suggested by Slingshot, setting "LP" as the root;
(D) Umap representation of the zoomed-in transitioning compartment used for pseudotime inference, colored according to pathway enrichment
scores.

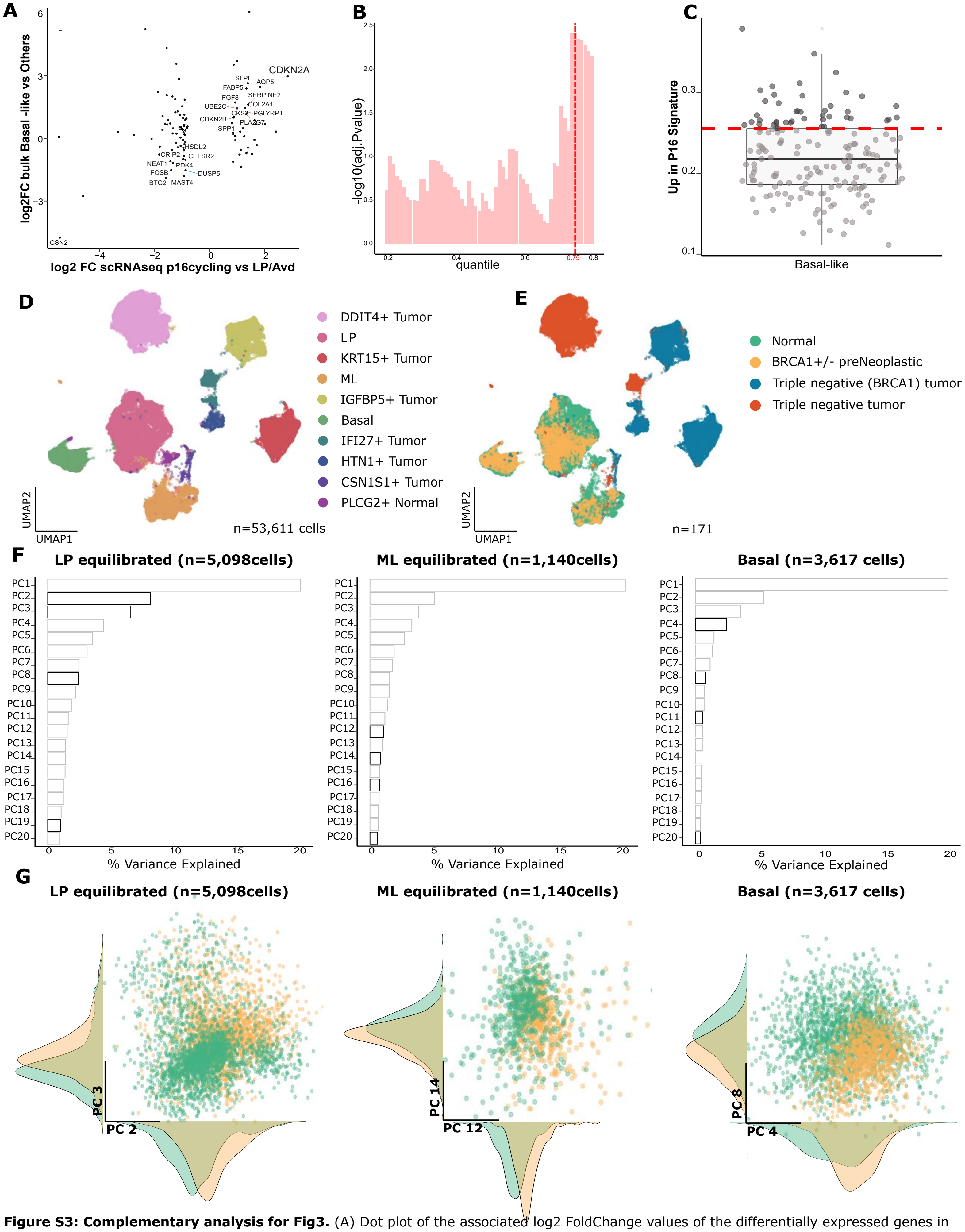


**Figure S3: Complementary analysis for Fig3.** (A) Dot plot of the associated log2 FoldChange values of the differentially expressed genes in the p16 cycling cells (from scRNAseq) and Basal-like samples (from bulk RNAseq);(B) Barplot representing the LogRank test adj.P-values according to the quantile of p16 signature discretization, red line corresponds to the 75th percentile (lowest P-value) (C) Boxplot representation of the p16 signature score within the basal-like samples from the PAN CANCER Breast dataset; the 75th percentile is indicated, which was used to discretize p16 signature values for the survival analysis presented in the figure. UMAP representation of the epithelial compartmentfrom the EMBO dataset, publicly available at GSE161529, where cells were colored according to their manually-annotated cell type (D) and sample of origin (E); (F) Barplot representation of the percentage of explained variance of the top 20 principal components (PCs) from PCA performed on each epithelial compartment, BRCA1 deficiency associated PCs were highlighted in black, (G) PCA representation of the two most informative PCs for each epithelial compartment, cells were colored according to heir sample of origin;

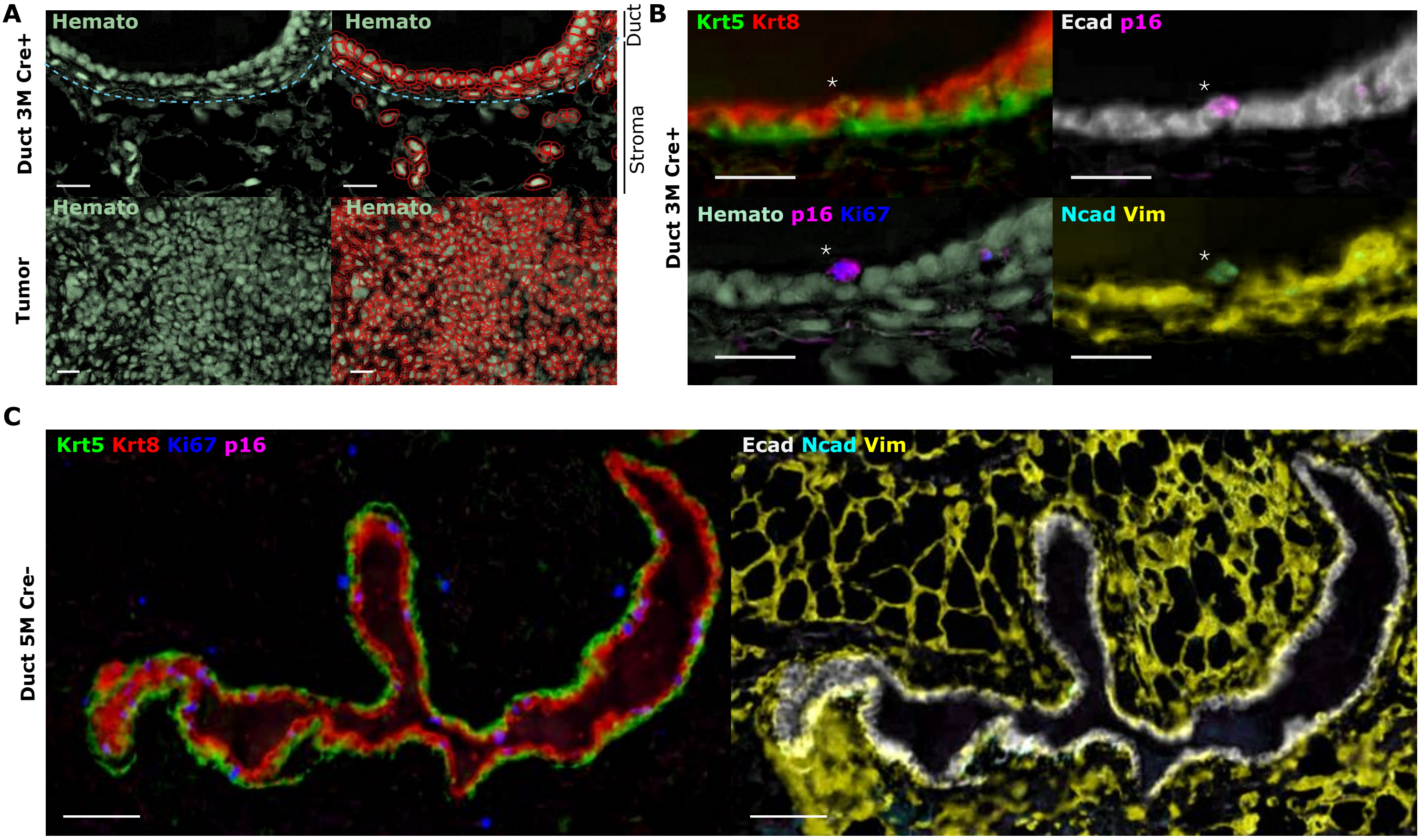


**Figure S4: Complementary analysis for Fig4.** (A) Example of automatic cell detection done on QuPath based on the hematoxylin staining for duct and tumor sample, scale bare represent 25µm. (B) Detail of the multiplex staining for all the markers, Hemato (light green), Cdkn2a (magenta), Ki67 (blue), Ecad (white), Krt5 (green), Krt8 (red), Ncad (cyan), Vim (yellow), scale bare represent 25µm. (C) Multiplex staining for a representative control duct from a 5 months Cre- mouse stained for Hemato (light green), Cdkn2a (magenta), Ki67 (blue), Ecad (white), Krt5 (green), Krt8 (red), Ncad (cyan), Vim (yellow), scale bare represent 50µm.

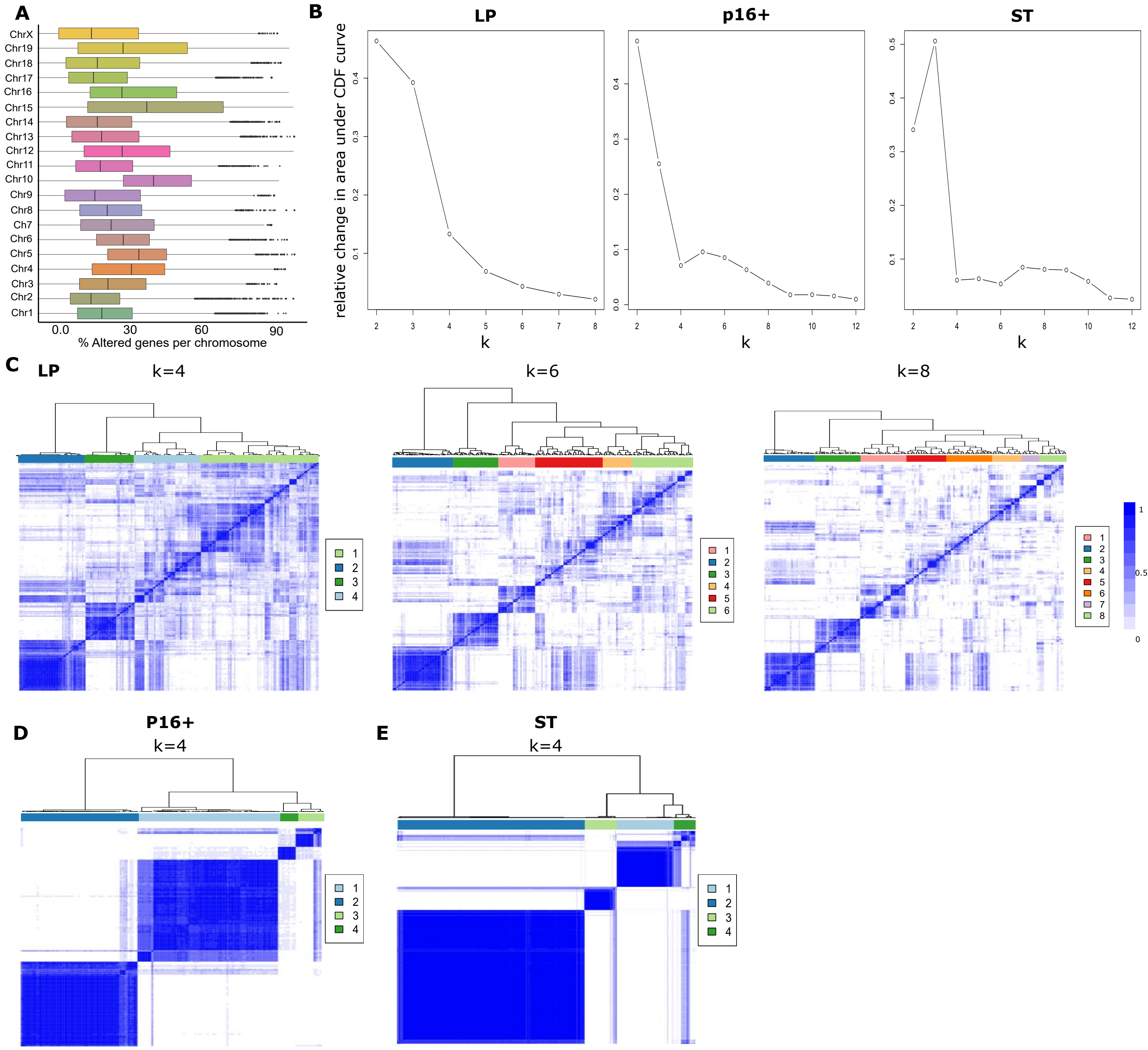


**Figure S5: Consensus clustering on CNV alterations.** (A) Boxplot distribution of the percentage of alterations in each chromosome within the epithelial compartment; (B) Distribution of the relative change in area under CPF values according to increasing numbers of clusters (k) for LP, P16+ and ST cells; (C) Heatmap representation of the corresponding consensus matrix to different values of the number of clusters (k=4,6,8) between LP cells; Heatmap representation of the corresponding consensus matrix to the optimal number of clusters (k=4) for both P16+ cells (D) and ST (E).

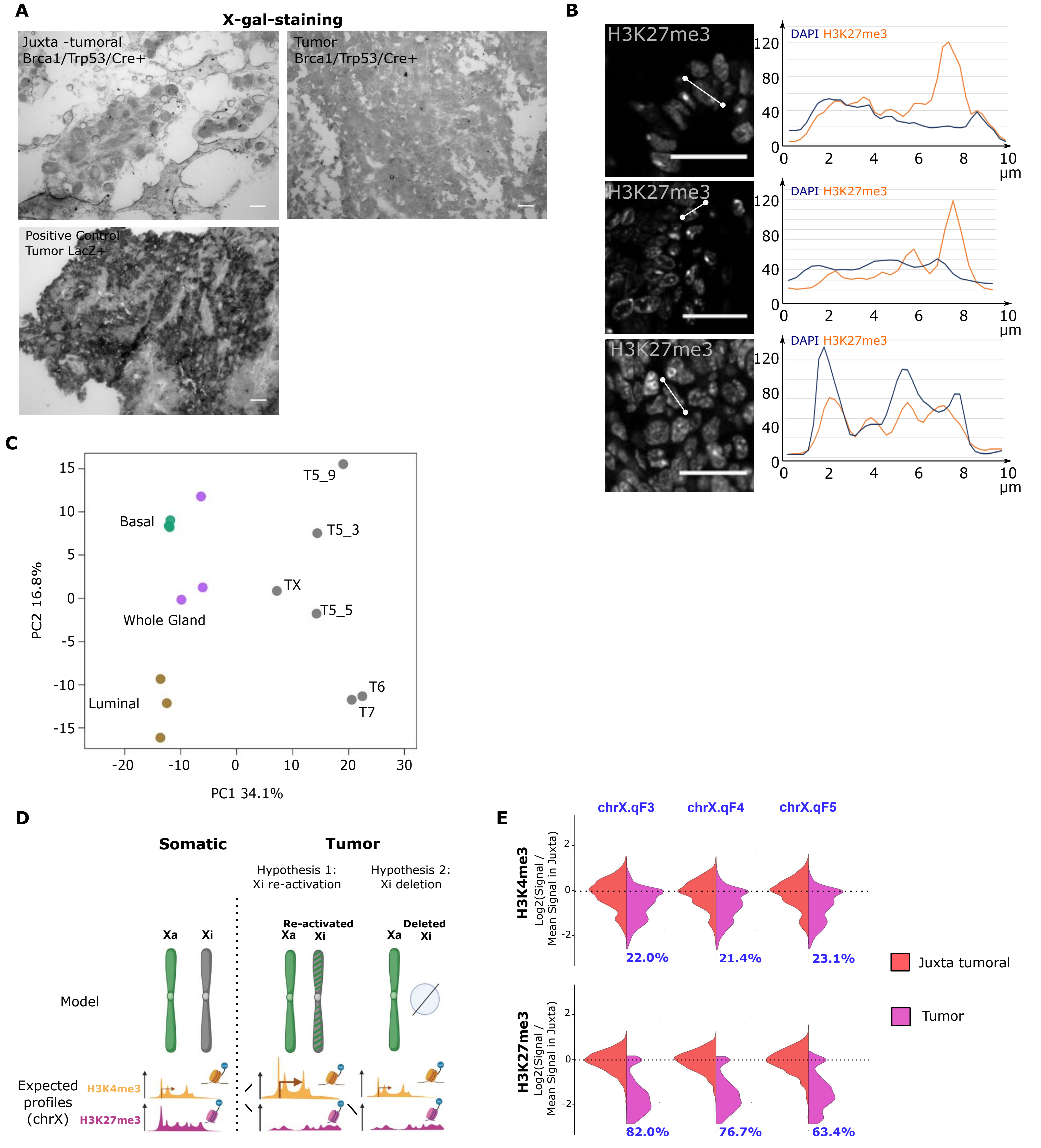


**Figure S6: Complementary analyses to Figure 6.** (A) Beta-Galactosidase (BGal) staining on multiple samples including, juxta-tumoral and Lacz +/- tumor tissue sections; (B) Representative sections of a mammary gland stained by immunofluorescence for H3K27me3, revealing the Xi and the micro Foci distributions; (C) PCA projection of normalized genome-wide H3K27me3 enrichments using peak-based annotation for tumor samples and sorted basal and luminal cell populations, on the first 2 principal components; (D) Simple scheme showing the hypothesis of the loss of inactive chromosome X that could fit the observations. Pointed arrows represent actively transcribed genes while flat-ended arrows represent repressed transcription. (E) Violin plots of the log-ratio of signal by the mean signal in juxta tumoral samples for 3 cytobands within the X chromosome for scH3K4me3 and scH3K27me3. Percentage tumor cells with gains or loss is shown above or below each violin.
